# Single-cell deconvolution of 3,000 post-mortem brain samples for eQTL and GWAS dissection in mental disorders

**DOI:** 10.1101/2021.01.21.426000

**Authors:** Yongjin Park, Liang He, Jose Davila-Velderrain, Lei Hou, Shahin Mohammadi, Hansruedi Mathys, Zhuyu Peng, David Bennett, Li-Huei Tsai, Manolis Kellis

## Abstract

Thousands of genetic variants acting in multiple cell types underlie complex disorders, yet most gene expression studies profile only bulk tissues, making it hard to resolve where genetic and non-genetic contributors act. This is particularly important for psychiatric and neurodegenerative disorders that impact multiple brain cell types with highly-distinct gene expression patterns and proportions. To address this challenge, we develop a new framework, SPLITR, that integrates single-nucleus and bulk RNA-seq data, enabling phenotype-aware deconvolution and correcting for systematic discrepancies between bulk and single-cell data. We deconvolved 3,387 post-mortem brain samples across 1,127 individuals and in multiple brain regions. We find that cell proportion varies across brain regions, individuals, disease status, and genotype, including genetic variants in TMEM106B that impact inhibitory neuron fraction and 4,757 cell-type-specific eQTLs. Our results demonstrate the power of jointly analyzing bulk and single-cell RNA-seq to provide insights into cell-type-specific mechanisms for complex brain disorders.

## Introduction

The progression of most neurodegenerative and neuropsychiatric disorders, including Alzheimer’s disease (AD), commonly disrupts a broad spectrum of regulatory networks at the genomic and epigenomic levels, and poses a significant challenge to elucidating the mechanisms underlying the disease progression. In the genome-wide association studies (GWAS), AD is highly heritable, and most of the heritability is explained by common genetic variants. However, it is also highly polygenic, involving potentially hundreds of independent regulatory mechanisms.^1–3^From the transcriptomic and epigenomic profiling of postmortem samples across different brain regions, we discover the target genes and regulatory elements perturbed by the disease progression and gain insights into the mechanisms of the genetic variants through the regulatory networks.^3–5^However, many different factors contribute to the variability of the existing postmortem brain data, and it is crucially important to identify and classify these factors by information source.

A large part of the complexity in mental health traits stems from cell-type heterogeneity in brain tissues.^6^Transcriptomic and epigenomic profiling at single cell-level resolution provides a principled tool with which to investigate the relationship between changes in cell-types and AD pathology. In our recent study, using single-cell RNA-seq (scRNA-seq) data across 80,660 nuclei isolated from post-mortem brain samples across 48 individuals, we discovered seven major types and 40 subtypes of the brain cells. We used these data to recognize cell-type-specific alterations associated with diverse pathological variables including age, sex, and AD pathology.^6^However, single-cell RNA-seq profiling is often limited to a small number of individuals and optimized to yield a large number of cells. However, most single-cell studies, including our published scRNA-seq analysis,^6^only involve at most tens of individuals:^7^too few to measure correlations with other population-level variables such as genotype information. At the bulk tissue-level, however, RNA-seq studies routinely profile hundreds of individuals. For example, the Religious Orders Study and the Memory and Aging (ROSMAP) study has profiled bulk RNA-seq from more than 400 individuals with matched genotype information^3,8^, and the Genotype-Tissue Expression consortium (GTEx) has profiled more than 2,500 brain samples across 13 brain regions from genotyped individuals.^9^

Here we develop a novel integrative framework to uncover cell-type-specific alterations of bulk samples, by combining both single-nucleus and bulk RNA-seq data with computational deconvolution followed by comprehensive association analysis. We develop a highly accurate deconvolution method which takes into account individual-level heterogeneity present in both single-nucleus and bulk data. We also directly address systematic discrepancies between single-nucleus and bulk data by characterizing substantial technical inconsistencies between them and developing a transformation approach to overcome them. We apply this method systematically across 3,387 samples to study the variation of neural cell-types across brain regions and their association with other variables measured in the bulk data. We then interrogate the mechanisms at the resolution of pathways and genetic regulatory networks by deconvolving the tissue-level eQTL models into cell-type-specific models.

## Results

### SPLITR deconvolution accounting for biological covariates and bulk-vs-single-cell differences

Existing deconvolution methods^10,11^estimate cell-type fractions from bulk RNA-seq data by making the explicit or implicit assumption that bulk RNA-seq data should match the sum of the same set of scRNA-seq data across the different cell types. In practice, however, aggregated scRNA-seq data and bulk RNA-seq data show substantial discrepancies, even for the most established marker genes. One reason for these discrepancies is that single-cell data and bulk data have highly distinct biases due to gene length, mRNA subcellular localization, transcriptional burstiness, mRNA stability, and the cell-to-cell variability of each gene’s expression patterns. This is most pronounced in single-nucleus RNA-seq datasets, as they only capture the nuclear component of each cell’s expression profile. Thus, aggregation of individual single-nucleus expression profiles is not expected to match bulk RNA-seq profiles that also capture the cytoplasmic component of each gene’s expression levels. Systematic corrections are therefore required to relate single-cell datasets into bulk datasets, which are currently not known.

In addition, existing deconvolution methods typically use a single reference profile for each cell type.^10,11^Such profiles are sometimes obtained by averaging multiple cells of the specific cell type,^12–14^and other times by using a predicted developmental trajectory.^15^However, biological variables such as disease status, age, or biological sex can substantially influence expressions of marker genes in both single-cell RNA-seq and bulk RNA-seq samples, making it inappropriate to use the same cell-type-specific reference expression profile for each individual. For example, the expression of several neuronal markers alters with age, sex, and disease status. Similarly, markers for nearly all cell types vary in their expression based on the phenotypic status of each individual. New methods are therefore needed which can tailor the cell-type-specific reference expression profiles used for each individual to their biological covariates.

To address these challenges, we developed a new deconvolution method, SPLITR (for Single-cell Phenotype-aware deconvoLution across Individuals from Total RNA-seq), which explicitly models: (1) inter-individual variation in both bulk and cell-type-specific gene expression levels across biological covariates including age, biological sex, and disease status; and (2) platform-specific biases and differences between single-cell and bulk RNA-seq datasets, including differences in subcellular localization of each gene’s mRNA population in the nucleus/cytoplasm. We achieve this by executing the following three steps of model estimation.

In step 1 of the SPLITR method (Fig. 1a), we use reference single-cell datasets to establish marker genes and reference average expression levels for each target cell type. Here, we focus on brain cell types and use snRNA-seq profiles that our group previously generated across 80,000 cells from 48 individuals, clustered into seven cell types, consisting of excitatory neurons, inhibitory neurons, astrocytes, oligodendrocytes, oligodendrocyte progenitor cells (OPCs), microglia, and pericytes & endothelial cells.^6^We used these clusters to define a set of 117 marker genes that were the most characteristic of each cell type, based on their differential cell-type-specific gene expression patterns (Supplementary Fig. 1), and confirmed these marker genes agreed with cell-sorted expression profiles^16^and independent single-cell expression profiles.

**Figure 1.**
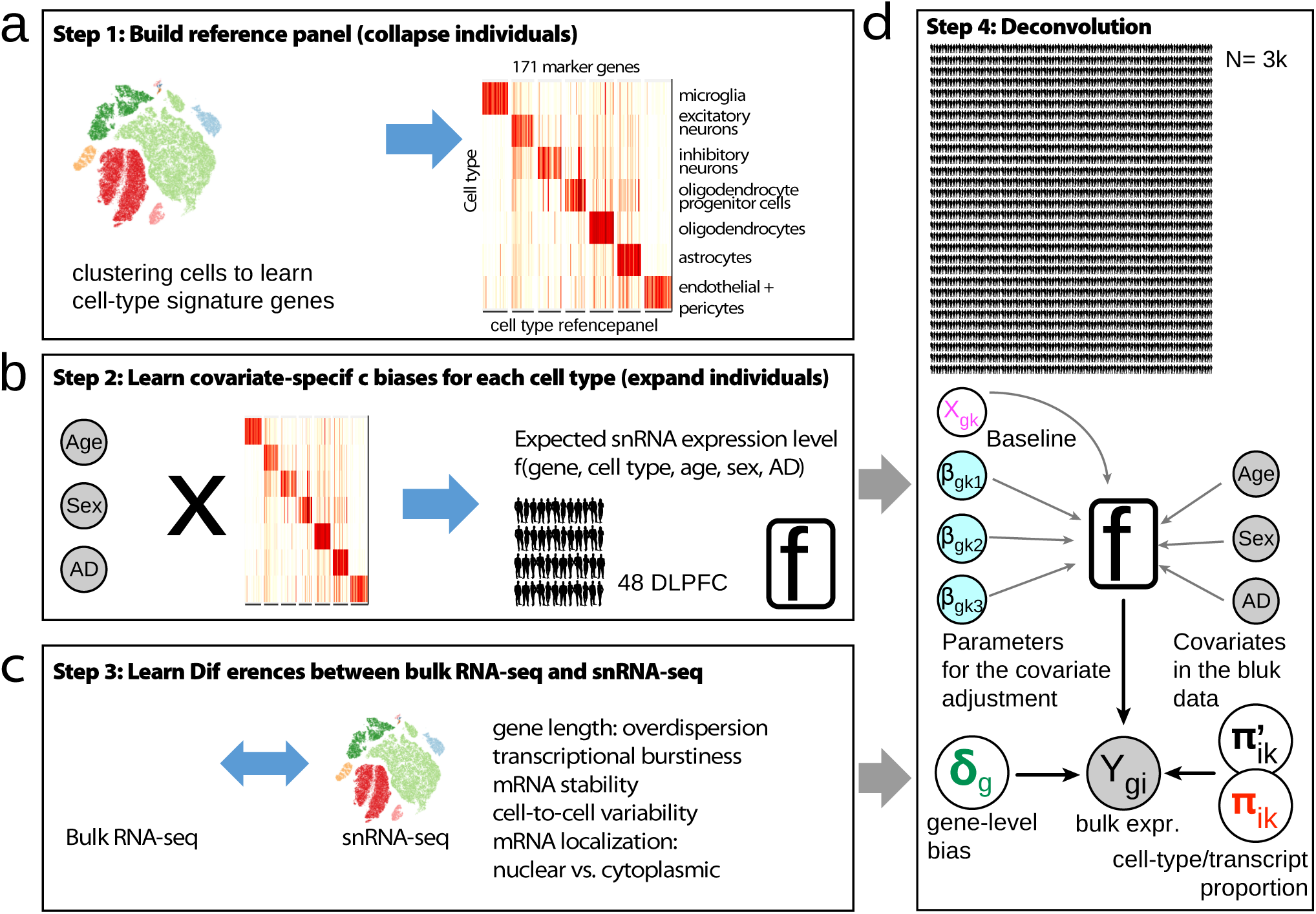
Overview of our deconvolution method.

In step 2 (Fig. 1b), we study the impacts of three phenotypic/biological covariates on cell-type-specific expression profiles. We use the cell-type-specific expression matrix (pseudo-bulk), created by taking average values over 80k cells across the 48 individuals within the seven cell types. We learn effects of the covariates by estimating a negative binomial model regressing cell-type-specific gene-vectors on the covariate and intercept terms. This modeling enables us to establish phenotype-aware adjustments to cell-type-specific expression patterns according to the phenotypic covariates of each individual whose bulk RNA-seq expression levels are to be deconvolved. For the application to Alzheimer’s Disease, we used age, sex, and pathological AD status, given their correlation with global gene expression changes in each major brain cell type.

In step 3 (Fig. 1c), we build a model that adjusts each marker gene’s bulk expression levels for experimental platform differences between scRNA-seq and bulk RNA-seq learn gene-specific correction terms. In particular, we find that scRNA-seq, and single-nucleus RNA-seq in particular, show gene-specific systematic differences stemming from each gene’s mRNA localization patterns, gene length, transcriptional burstiness, mRNA stability, cell-to-cell variability, and nuclear vs. cytoplasmic fractions. We assume the overall impact of such biases is shared across individuals (samples), and estimate the corresponding terms by leveraging control genes that show consistent expression patterns between the bulk and snRNA-seq data (see Online Methods for details).

In step 4 (Fig. 1d), we use these learned parameters to deconvolve the bulk expression profile by fitting a negative binomial regression of the bulk profile, adjusting for the learned phenotype-specific correction terms for each individual, and the platform-specific correction terms for each gene. For each cell type, we calculate an activity fraction estimate, corresponding to the number of transcripts produced by each cell, and a cell proportion estimate, correcting for the overall activity level of each cell type (Supplementary Fig. 3). In the case of brain, for example, we found that neuronal cells generate nearly four times more mRNA transcripts than glial cells, so their activity fraction estimates are approximately four times larger than their cell proportion estimates.

### Deconvolution of 3,386 bulk samples and experimental validation of cell type proportions

We used SPLITR to deconvolve a total of 3,386 bulk RNA-seq post-mortem brain samples, encompassing 15 brain regions from 1,127 individuals across three different studies (Fig. 2): (1) 482 dorsolateral prefrontal cortex (DLPFC) samples from the Religious Orders Study and Memory and Aging Project (ROSMAP) from Rush University^3,17^; (2) 263 temporal cortex samples from the Mayo Clinic Brain Bank^18^, and (3) 2,642 samples across 13 brain regions in the Genotype-Tissue Expression (GTEx) data^9^.

**Figure 2.**
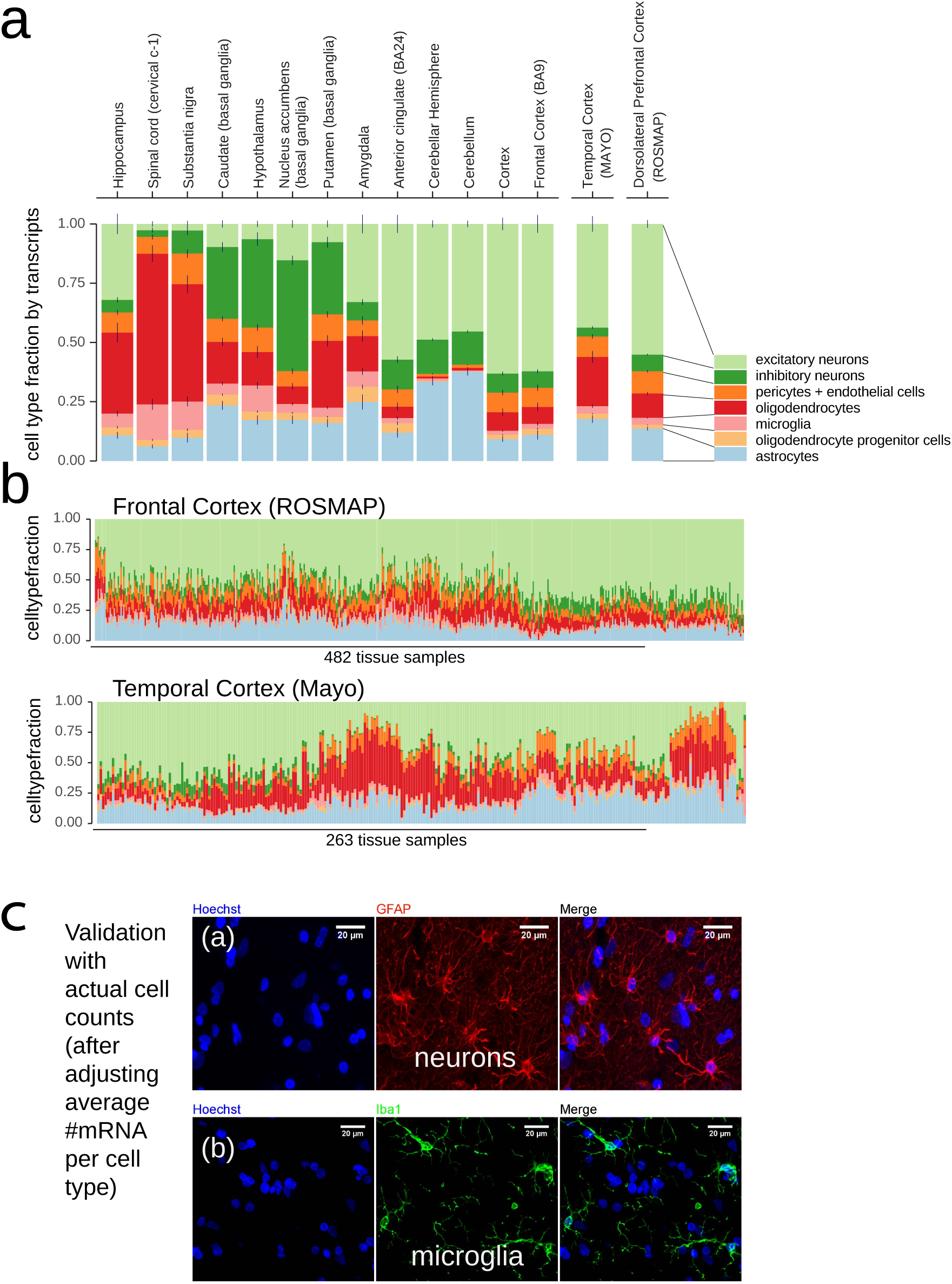
Deconvolution of 3,386 tissue-level RNA-seq data across 15 brain regions reveals unique cell type compositions across different brain regions. Different colors indicate different cell types. (a) Average cell type fractions across different brain regions. (b) Population-level variation of cell type compositions in ROSMAP and Mayo cohorts. (c) The estimated cell type compositions are experimentally validated against the cell counts in microscopic images of neurons and microglial cells.

Despite common discrepancies in the literature between different methods of estimating cellular proportions in brain samples^19^, we found that the resulting activity-corrected cell type fraction estimates were consistent with several previously-reported fractions from direct measurements. For example, the average cell count fractions estimated from the 482 dorsolateral prefrontal cortex samples in the ROSMAP cohort were 31% for neurons, 51% for glial cells, and 18% for vascular cells, consistent with previous estimates using the fractionator sampling method^20^and the isotropic fractionator^21^. Excitatory neurons were 4 times as abundant as inhibitory neurons, in line with previous reports^22^.

We also experimentally confirmed these cell type fractions using immunostaining in matched samples (Fig. 2c). Our SPLITR estimated of astrocytes were 18% of cells (±5%), consistent with our immunostaining-measured average of 18% across 8 ROSMAP samples. Even for microglia, which are both less abundant and smaller cells, thus biasing their abundance in some single-cell preparations, our SPLITR-based estimate of 9% (±5%) was consistent with our immunostaining-measured average of 10%.

The cell type proportion estimates sometimes differed from the counts of nuclei obtained for each cell type from the 10X protocol^6^, with the single-nucleus counts sometimes closer to our activity fraction estimates. This is likely due to experimental biases in earlier versions of the 10X protocol to more efficiently capture larger nuclei with more transcripts. For example, astrocytes, microglia, pericytes, and endothelial cells were under-represented in our 10X datasets, while excitatory neurons and oligodendrocytes were overrepresented compared to previous estimates. This indicates that SPLITR deconvolution can provide accurate estimates of cell type proportions, even when based on single-cell datasets with skewed proportions, as it is based on the expression patterns inferred from the single-cell datasets, rather than the cell proportions in those datasets.

Our estimated cell type proportions also captured known variability across different brain regions. For example, we found a substantially higher fraction of oligodendrocytes in hippocampus than frontal cortex (48% vs. 16% on average), more microglial cells in hippocampus than cortex (12% vs. 6% on average), and fewer neurons in hippocampus than in temporal cortex or in frontal cortex (15% vs. 21% vs. 42% on average).

Our estimates also captured cell type proportions vary across different cohorts associated with the different age ranges of the individuals profiled (Supplementary Tab. 1). Comparing the younger GTEx cohort (59 years old on average) with the older ROSMAP cohort (88 years old on average), we found that neurons decreased from 42% to 31% of cells, while glia increased from 38% to 50%, consistent with neuronal loss during aging.

We also compared the results of SPLITR with CIBERSORT^10^, using the same marker gene profiles (Supplementary Fig. 2). We found a general agreement for higher-abundance cell types, including excitatory neurons, oligodendrocytes, astrocytes, and microglia. However, for two of the cell types, CIBERSORT showed systematic problems, resulting in 0% estimates for inhibitory neurons for 68% of samples, and 0% estimate for oligodendrocyte progenitor cells for 79% of samples. Moreover, while SPLITR captured differences due to sex and age, CIBERSORT did not capture these subtle differences (Supplementary Fig. 2b-c).

### Discovery of genetic variants influencing cell type proportion

We first sought to recognize genetic variants that may underlie these cell type proportion differences between individuals. Treating the SPLITR-inferred proportions of each of the 7 cell types as a quantitative trait, we carried out a cell-fraction genome-wide association study (cfGWAS) to recognize genome-wide significant and sub-threshold loci associated with cell type fraction (cfGWAS hits), which we define as single-nucleotide polymorphisms (SNPs) that govern cell type proportions. Using 403 ROSMAP individuals that have both genotype and RNA-seq data, we found several genome-wide significant (P<5e-8) and sub-threshold (P<1e-5) associations with cell type proportions.

The strongest association (P=6.4e-09) was between reduced excitatory neuron fraction and >100 SNPs in the *TMEM106B* locus of chromosomal segment 7p21.3, including the A-to-G rs1990620 SNP (Fig. 3a). This locus is not previously-associated with AD, but it is associated with an AD-related neurodegenerative disease, frontotemporal lobar degeneration with TDP-43 inclusions (FTLD-TDP), and also with decreased-neuronal-fraction allele showing increased FTLD-TDP risk and decreased cognition in amyotrophic lateral sclerosis (ALS)^23–26^. Indeed, this association was not due to AD pathology in our cohort, and the cell-type-proportion association remained statistically significant after conditioning on the pathological phenotypic variables, such as the accumulation amyloid-beta and neurofibrillary tau (NFT) proteins, and pathogical AD. These results suggest that the *TMEM106B* locus is an AD-independent contributor to cognitive decline, via decreased neuronal fraction.

**Figure 3.**
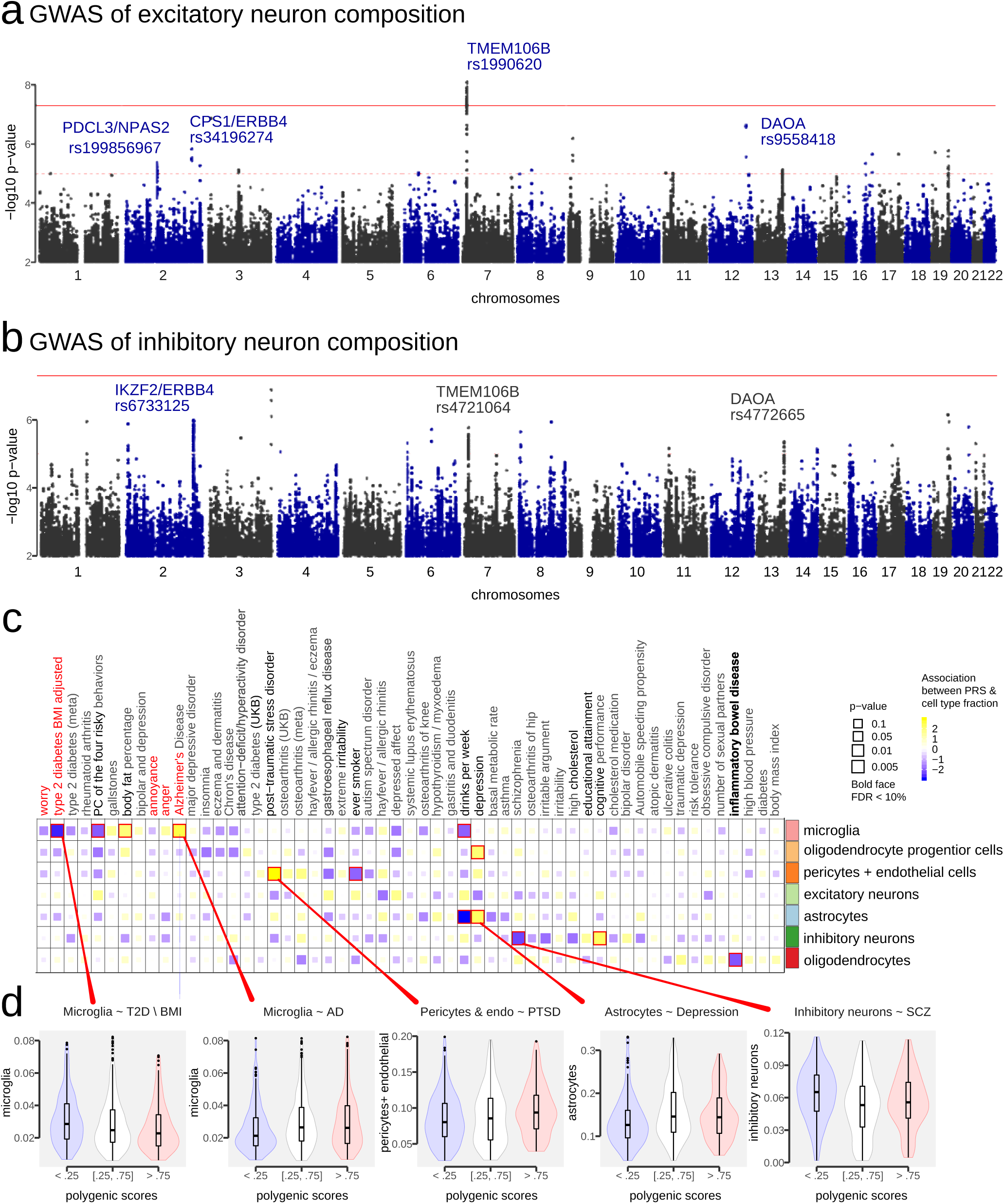
Genetic variation across individuals induce markedly differential cell type compositions. (a) Genome-wide association studies of excitatory neuron fraction. X-axis: genomic location across 22 autosomes; y-axis: -log10 p-value of association effect sizes. (b) Genome-wide association studies of inhibitory neuron fraction. (c) Association of cell type fractions with the polygenic risk scores of 56 relevant traits. The boxes are scaled proportionally to association p-value (larger, more significant). The colors reflect the directionality of associations (blue for negative, yellow for positive). (d) Examples of the significant associations between cell types and GWAS traits at the polygenic risk score-level. X-axis: quantile of polygenic scores; y-axis: cell type fractions estimated from ROSMAP cohort.

Genotype-associated expression variation indicates that two nearby genes may mediate the *TMEM106B* locus genetic effect on neuronal fraction: the *TMEM106B* gene itself, a transmembrane gene invoved in dendrite morphogensis and the regulation of lysosomal trafficking, and *GRN*, an age-associated^27^essential gene involved in tau-negative FTLD^28^and lysosomal dysfunction during FTLD progression.^29^Both genes showed significantly-reduced tissue-level expression associated with the decreased-neuronal-fraction allele (p<3e-06 and p<2e-03, respectively), and previous studies indicate that *TMEM106B* interacts with *GRN*, and that rs1990620 may be the causal variant in this locus, via disruption of a CTCF binding motif that alters a topologically-associated domain and up-regulates *TMEM106B*^30^. The associations were still significant when only including the controls (p<9e-04 and p<1.5e-02, respectively), and cell-type-specific eQTL analysis with cell-sorted and snRNA-seq data confirmed over-expression of *TMEM106B* in neurons, astrocytes, and oligodendrocytes, consistent with the previous reports^31,32^.

Several additional subthreshold-level associations with excitatory neuron fraction were found overlapping known causal genes in neurodevelopmental processes, including: *ERBB4*, a risk gene for schizophrenia and a selective and functional marker gene for glutamatergic^33^and GABAergic synapses^34^in inhibitory neurons and interneurons; *DAOA* associated with schizophrenia in an Asian cohort^35^; and *NPAS2*, conferring neuropsychiatric anxiety disorder and regulating GABAergic signal transmissions^36^. While the main signal in the *TMEM106B* locus affects excitatory neurons, we found an additional genetic signal associated with both inhibitory neurons associated with rs1990620 (P=3.08e-6). Lastly, we found an additional association with inhibitory neurons within the *TMEM106B* locus, with cfGWAS SNP rs4721064 (p=1.69e-06), whose effect is independent and additive with that of rs1990620.

### Deconvolved cell-type fractions are associated with increased risk for diverse phenotypes

We found that changes in cell type fraction were also associated with increased risk for multiple traits, even when these were not directly measured in our cohort (Fig.3c-d). For all 944 genotyped individuals across ROSMAP and GTEx, we used their complete genotype information across millions of common variants to calculate their genetic risk for a set of 56 traits (Supplementary Tab. 2), using polygenic risk score (PRS) estimates from GWAS summary statistics data (p-value threshold 0.01, with linkage disequilibrium decorrelation^37,38^instead of pruning). A total of 17 traits showed nominal significance (p-value<0.05), including Alzheimer’s Disease and Crohn’s disease.

We found several noteworthy examples (Fig. 3c-d): higher microglial fraction was associated with increased AD risk and increased body fat percentage, but decreased risk for type 2 diabetes (T2D) in ROSMAP; higher oligodendrocyte progenitor cell fraction was associated with increased risk of depression; higher pericyte and endothelial fractions was associated with increased risk of post-traumatic stress disorder (PTSD) and decreased risk of smoking; higher astrocyte fraction was associated with increased risk of depression and decreased risk of drinking; higher inhibitory neuron fraction was associated with higher cognitive performance and lower schizophrenia risk; lastly, an increased oligodendrocyte fraction was associated with decreased risk of Inflammatory Bowl Disease (IBD).

Many traits associated with the same cell type showed only negligible correlation at the overall PRS level, indicating that our method can capture correlations not directly visible using only genetic information.

### Cell-type fraction differences associated with Alzheimer’s pathology, biological sex, and age

We next investigated whether cell type proportion changes were associated with phenotypic differences between individuals within the ROSMAP cohort, where phenotypic variables are readily available (Fig. 4).

**Figure 4.**
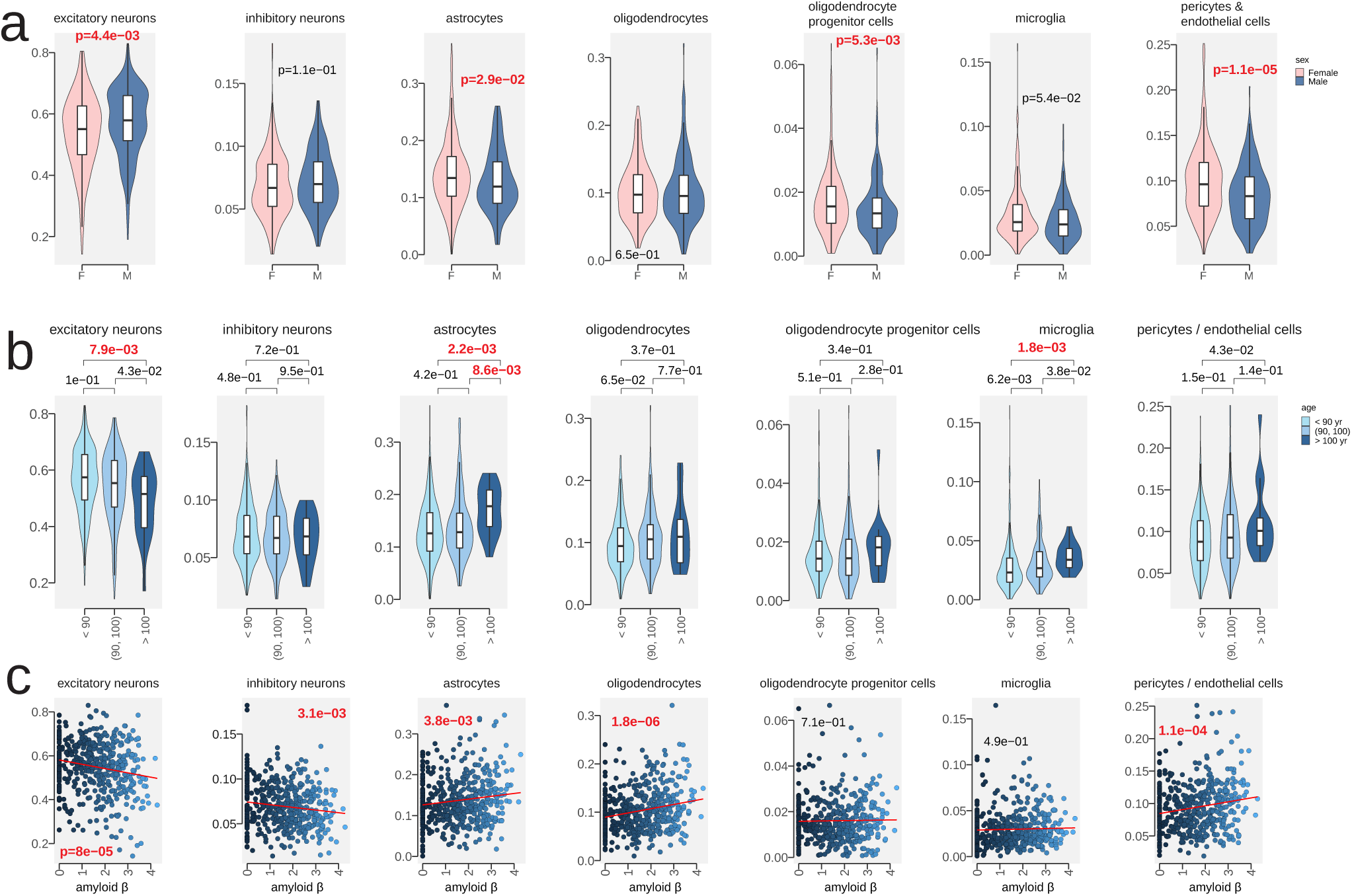
Cell-type fractions in the brain samples dramatically change with known pathology and demographic variables. Y-axis: cell type composition; X-axis: pathological and demographic variables. (a) Cell type compositions change between female and male samples. X-axis: female vs male. (b) Cell type composition changes across different age groups. X-axis: age groups (years at death). (c) Cell type compositions as a function of pathological variables. X-axis: amyloid-beta (in square root).

We found that AD-related pathological variables were strongly associated with cell type proportion differences (Fig. 4c; Supplementary Fig. 4). Amyloid-beta deposition showed the strongest associations with fewer excitatory neurons (P<8e-5), fewer inhibitory neurons (P<3e-3), more oligodendrocytes (P<2e-6), more astrocytes (P<3.8e-3), and more pericytes/endothelial cells (P<1e-4). Tau-protein deposition and loss of cognition also showed significant associations with fewer excitatory neurons and more oligodendrocytes.

We also found that cell type proportions were strongly associated with both biological sex and age (Fig. 4a-b). Male samples showed a higher fraction of excitatory neurons than female samples (P<0.004, Wilcoxon rank-sum test) and a lower fraction of astrocytes (P<0.03), oligodendrocyte progenitor cells (P<.006), and vascular cells (P<2e-5) (Fig. 4a). In addition, older individuals (>100 years old) showed different cell type proportions than younger groups (<90 years old), with fewer excitatory neurons (P<0.008), more astrocytes (P<0.003), and fewer microglia (P<0.002) (Fig. 4b).

These results indicate that our deconvolved cell type fractions successfully capture cell type proportion changes associated with phenotypic differences, even though only bulk samples were utilized for these analyses.

### Cell-type-specific gene expression changes in AD show biologically-meaningful functional enrichments

We found 2,470 genes with cell-type-specific changes associated with amyloid-beta, neurofibrillary tangles, and episodic memory decline in one of the seven cell types (Fig. 5a-b), using a generative model that captures the relationships between each gene’s transcript level with an interaction term of cell type and each pathological variable (age, sex, RIN scores, and other phenotypes). We controlled the FDR at 4.4% with the null distribution constructed by the Freedman-Lane permutation^39^of only one interaction term at a time while fixing all other correlated variables (Fig. 5a, Supplementary Fig. 5a). Only 12 of these 2,470 genes are among the 171 cell-type-marker genes.

**Figure 5.**
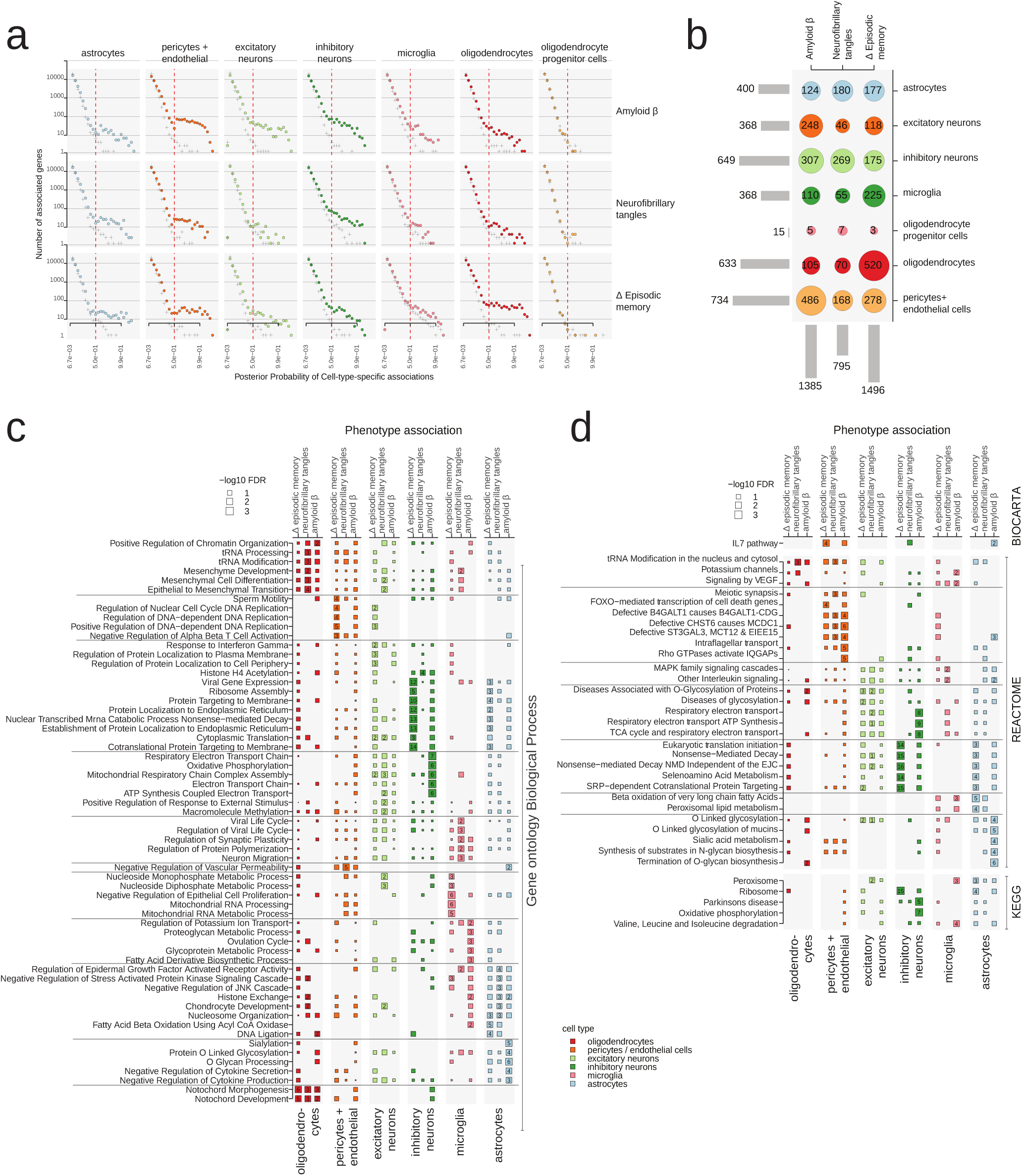
Cell-type-specific gene expressing chanes in AD phenotypes show biologically-meaningful functional enrichment. (a) We estimated the null distribution (y-axis) for the posterior probability (x-axis) of non-zero cell-type-specific gene-level associations by Freedman-Lane permutation^39^(marked by “+”). Using them, we empirically calibrated the false discovery rate of the optimized posterior probabilities of genes (marked by solid dots). (b) Controlling FDR at 4.4% (posterior probability > 0.5), we found 2,470 genes are significantly associated in a cell-type-specific manner. The number in each circle accounts for the genes found associated with a different phenotype (the column) in a particular cell type. (c) Cell-type-specific gene ontology (biological process) enrichment results for the significantly-associated genes. X-asis: cell types; y-axis: keywords (gene sets). (d) Cell-type-specific pathway enrichment results for the significantly-associated genes.

These 2,470 genes showed highly cell-type-specific enrichment across 191 gene ontology (GO) terms (Fig.5c) and 88 MSigDB^40^canonical pathways (Fig.5d) (FDR < 5%). Distinct enrichments were sometimes found for distinct AD phenotypes, between memory loss, neurofibrillary tangles, and amyloid-beta.

For example, genes with inhibitory-neuron-specific expression differences associated with AD pathology were enriched in intracellular transport (including endoplasmic reticulum) for memory-associated expression changes, and with mitochondrial biology for amyloid-associated changes. Genes with oligodendrocyte-specific expression differences associated with AD pathology were enriched in notochord development for memory-associated changes consistent with their roles in remyelination^41,42^and with our single-cell analysis results^6^, and with mesenchymal differentiation for tangles-associated changes. Genes with microglia-specific expression differences in AD were enriched in synaptic plasticity^43^for neurofibrillary-tangles-associated expression changes, in mitochondrial functions for memory-associated expression changes, and fatty acid metabolism for amyloid-beta-associated expression changes. Genes with astrocyte-specific expression differences in AD were enriched in cytokines and secretion for amyloid-beta-related phenotypes, consistent with secretion of pro-inflammatory cytokines in astrocytes with the accumulation of amyloid-beta^44^.

These results reveal a complex set of cell-type-specific alterations in diverse pathways associated with distinct phenotypic signatures of AD, provide important insights into the cellular and molecular changes in AD, and demonstrate SPLITR’s ability to recognize cell-type-specific from bulk RNA expression.

### Sparse Bayesian regression deconvolves tissue-level genetic effects into cell-type-specific eQTLs

To help elucidate causal paths between genetic variation and complex brain disorders, we next sought to recognize genetic variants with cell-type-specific effects on brain gene expression, both at the bulk level and at the cell-type-specific level. For tissue-level eQTLs, we used our previously-described sparse Bayesian multivariate model^45^, and for cell-type-level eQTLs we developed a new Bayesian eQTL deconvolution framework that models the observed bulk genetic effects as a mixture of cell-type-specific genetic effects, and infers a cell-type-specificity score (between 0 and 1) for each eQTL gene (eGene) in each cell type, corresponding to the probability with which this gene has cell-type-specific genetic effects for that cell type (Fig. 6a; Methods). To compare the performance of our deconvolved multivariate approach, termed deQTL, with other interaction QTL methods, we simulated realistic gene expression data, embedding a single causal cell type for each gene. We repeated our experiments on 121 randomly-selected linkage disequilibrium (LD) blocks, varying the level of expression heritability and number of causal eQTL variants (see Methods). In power analysis, our proposed approach clearly outperforms the other methods frequently used in cell-type interaction QTL analysis (Fig. 6b). Moreover, under the high heritability regime (> 10%), the posterior probability of the deQTL model accurately distinguish causal cell types from the non-causal ones (Fig. 6c).

**Figure 6.**
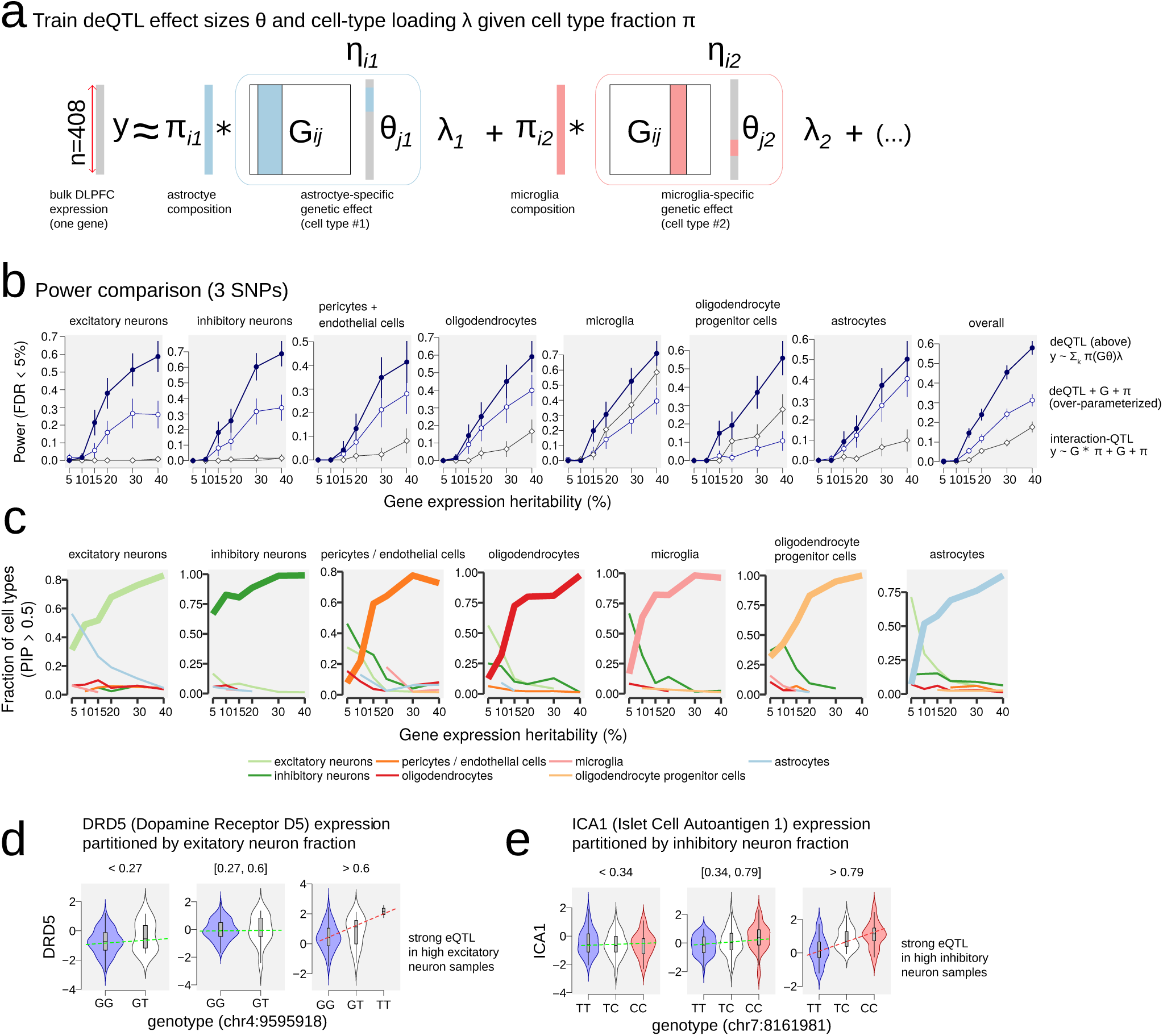
Deconvolved eQTL (deQTL) analysis dissects tissue-level genetic effects into the cell-type level mechanisms. (a) A schematic diagram for our deQTL method. Π: cell type fraction estimated by the deconvolution step. G: genotype matrix. *θ*: cell-type-specific multivariate effect size. *λ*: cell-type loading. *η*: cell-type-specific genetic effect aggregated over multiple SNPs. (b) Power comparison with competing methods. Different colors and shapes mark different methods. X-axis: gene expression heritability in simulation. Y-axis: statistical power of causal cell type identification with empirical false discovery rate (FDR) controlled at 5%. (c) Our deQTL method separates causal cell types (think lines) from non-causal ones (thin lines). Each panel, we simulate data assuming one causal cell type (the title of each panel). X-axis: gene expression heritability in simulation. Y-axis: the fraction of cell types discovered by the PIP cutoff > 0.5. (d-e) Examples of the genes that are significantly regulated in the deQTL models, but fail to reach significance under a marginal eQTL model. X-axis: genotype of the lead SNP; y-axis: quantile-normalized gene expression. Total ROSMAP samples are partitioned by the cell type fractions. (d) A significant deQTL on *DRD5*. (e) A significant deQTL on *ICA1*.

We applied this method to the 403 ROSMAP individuals that have both genotype information and gene expression information available. At the tissue level, we found a total of 5,586 eQTL genes (eGenes) with highly-heritable gene expression, associated with a total of 7,783 independent SNPs. At the cell-type-level, we found a total of 3,869 eGenes with cell-type-specificity score >0.9, associated with 4,757 independent SNPs. Approximately half of tissue-level eGenes (N=2,687, 48%) were also discovered at the cell-type level (Supplementary Fig. 5), enabling us to partition their genetic effects into the cell-types where they act.

A large fraction of cell-type-specific eGenes (N=1,182, 30%) were not discovered in our tissue-level analysis, indicating that our approach can discover high-confidence cell-type-specific eGenes even when these are not visible at the tissue level (Supplementary Tab. 3). For example, DRD5 showed no genetic association at the tissue level, but individuals with the TT allele of rs6448858 (chr4:9595918) were in the top 40% of samples with highest excitatory neuron content, resulting in a high interaction term in our model, and a high cell-type-specificity score (Fig. 6d). Similarly, ICA1 showed no tissue-level genetic effect, but individuals carrying the CC allele of rs6965329 (chr7:8161981) lay were among the 20% of samples with highest inhibitory neuron fractions (Fig. 6e).

Most cell-type-specific eGenes act in a single cell type (N=3,133, 81%), and a minority act in multiple cell types (N=736, 19%). Most act in inhibitory and excitatory neurons (61%), followed by oligodendrocytes (n=710), astrocytes (n=588), microglia (n=364), pericytes & endothelial cells (n=319), and oligodendrocyte progenitor cells (n=24) (Supplementary Tab. 1). For 872 cell-type-specific eGenes we found multiple independent eQTL variants, indicating more complex genetic control. Conversely, for 267 cell-type-specific eQTLs, we found multiple target eGenes, implicating gene-level pleiotropy.

### Stratification of the GWAS polygenic risk score (PRS) models by the deconvolved eQTL annotations

Lastly, we sought to recognize the cell types where disease-associated genetic variants exert their effect for diverse brain disorders, using genome-wide statistics for 56 neuronal, behavioral, psychiatric, and neurodegenerative traits (Supplementary Tab. 1). For each of the seven major cell types, we computed a PRS for each of the traits, using all nominally-significant (P-value<1e-2) SNPs that lie within a ±1 kb window of an annotated cell-type-specific eQTL for that cell type (Fig. 7a). We then calculated the enrichment for each cell type by comparing the cell-type-specific PRS score to the PRS score obtained using all the SNPs.

**Figure 7.**
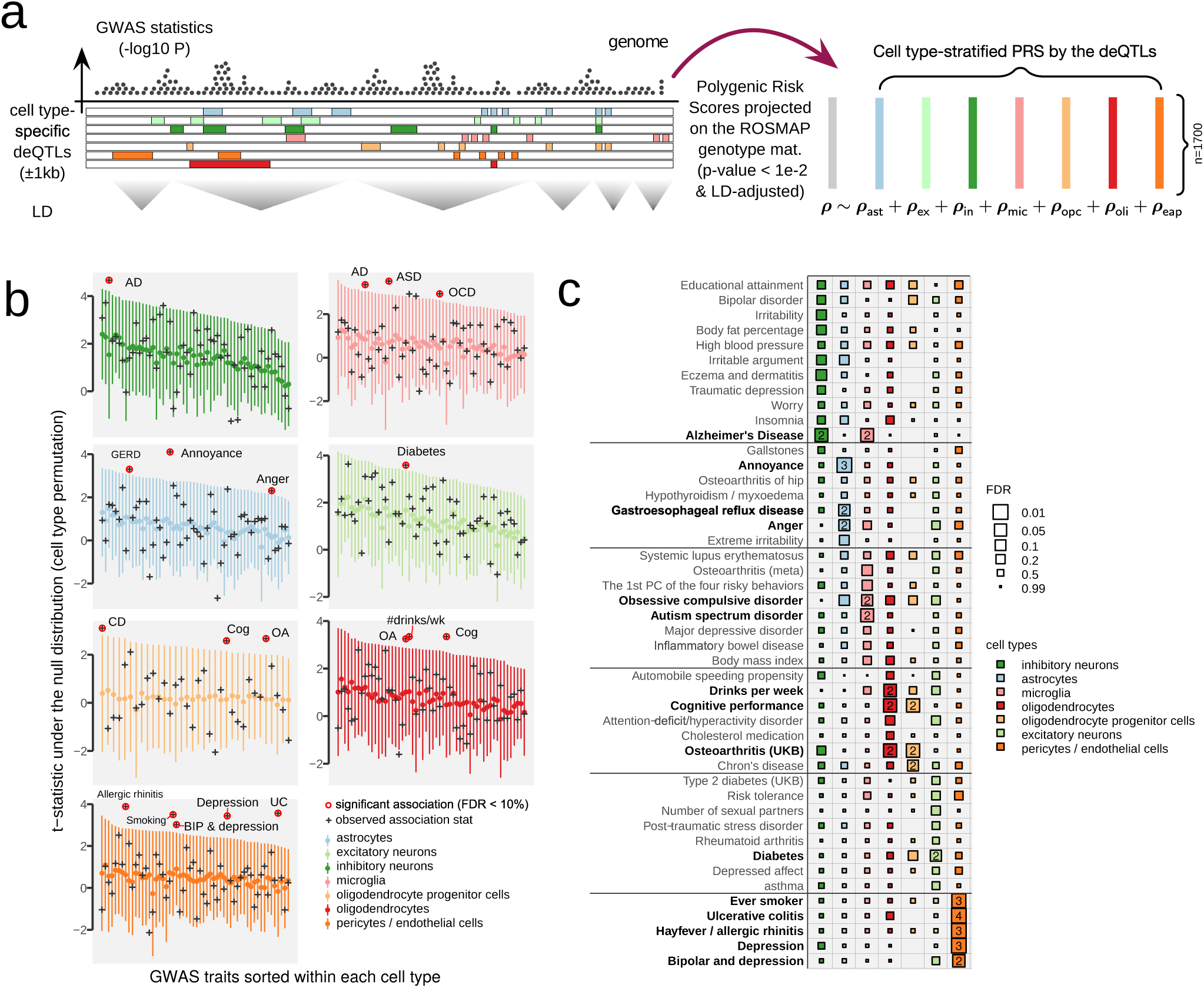
Deconvolved eQTL SNPs partition heritability of GWAS summary statistics into 7 cell types. (a)A schematic diagram showing our approach to partition GWAS data by significant deQTL SNPs. We construct 7 PRS models, each of which is stratified by the cell-type-specific genomic regions. (b)We calibrate the significance of cell-type-specific enrichment by cell-type label permutation. (c)Summary of significant cell-type-specific deQTL SNPs. The boxes are scaled proportionally to association FDR (larger, more significant). The colors indicate different cell types. For those significant enrichments (FDR < 10%), we denote the percentage of genetic variance explained by deQTLs.

Across all 1,682 individuals in the ROSMAP cohort^46^, we found 15 GWAS traits that show significant cell-type-specific PRS scores across 19 cell types (FDR<10%), indicating that genetic variants in that trait preferentially act through cell-type-specific eQTLs in that cell type (Fig. 7b-c). For example, we find that microglial-specific eQTLs contribute disproportionately to the risk scores of AD^2^, OCD (obsessive-compulsive disorder)^47,48^and ASD (autism spectrum disorder)^49^. Similarly, oligodendrocyte-specific eQTLs significantly enrich GWAS signals of osteoarthritis^50^and cognitive performance^51^. Pericyte and endothelial-specific eQTLs contribute disproportionately to increased risk of smoking^52^, UC (ulcerative colitis)^53^, allergy^54^, and depression and bipolar disorders^54^.

The importance of microglial cells in AD is well-recognized^55,56^, and several AD genes, such as BIN1^57^and MS4A4^58^, are shown to act specifically in microglial cells. For ASD, the previous analysis showed male-specific over-expression of microglial marker genes in the cortex^59^; for OCD, a mouse study showed that over-expression of NFKB/TNF-alpha pathways causally acts on relevant traits, such as excessive self-grooming behavior and hyperexcitability of the corresponding neurons^60^.

## Discussion

Understanding the mechanism of complex traits, including neurodegenerative disorders, has become a crucial component of prevention and treatment, yet remains a challenging and open problem. Part of the challenge stems from the complexity of the diseases at the cellular and molecular levels. A causal mechanism of complex traits is often manifested through multiple layers of genomic and epigenomic regulatory networks. The emerging technology of single-cell and single-nucleus sequencing provides unbiased profiling of cell types from a mixture of samples. Knowing the relevant cell-type context is a crucial step toward dissecting the complexity of diseases. Cell-type information enables biologically-informed Bayesian and causal inference, and improves experimental design in a matched cellular environment.

However, most single-cell-resolution profiling experiments cover a limited sample size and do not include the investigation of variation across individuals. On the other hand, while tissue-level bulk RNA-seq data fail to reach a cell-type resolution, they often carry a sufficiently large sample size. From richly-phenotyped bulk data, we can identify population-level associations of transcript measurements with other variables, such as genetic variants and phenotypes. Associations with small-effect variables are only made possible with a large cohort. Computational deconvolution methods, including SPLITR, abridge the gap between snRNA-seq and bulk RNA-seq data. We learn cell-type models from snRNA-seq and estimate cell-type fractions in the bulk data so that subsequent analysis can leverage a large sample size and rich phenotypic information.

Here, we present a highly calibrated deconvolution method, SPLITR, followed by a series of integrative studies with the variables in large-sampled bulk data. We identified cell-type-specific mechanisms of AD and other relevant disorders at the phenotype, demographic information, pathway, and genotype-level. Moreover, we characterized putative mechanisms, which may have impacted AD and other diseases, while pinpointing a molecular and pathway-level basis for understanding the comorbidity of complex neurodegenerative disorders. For instance, our results already suggest that microglial cells are a converging point of AD and neuropsychiatric disorders, such as OCD and ASD. Genetic markers in *TMEM106B* implicate potential pleiotropy between AD and FTLD in neuronal cells. Applying the same principle, we can investigate other neurodegenerative and neuropsychiatric disorders and even diseases in other domains, such as diabetes and cardiovascular disorders.

## Materials and Methods

### Preprocessing of the ROSMAP, Mayo, and GTEx RNA-seq data

We downloaded the ROSMAP RNA-seq data in the Dorsolateral Prefrontal Cortex (DLPFC) from Synapse (https://www.synapse.org/#!Synapse:syn3388564). We used gene-level expression data quantified by RSEM^61^, including a total of 55,889 coding and non-coding genes according to the GENCODE annotations (v19). The RNA-seq raw count data in the temporal cortex of 263 individuals from the Mayo RNA-seq project was downloaded from Synapse (https://www.synapse.org/#!Synapse:syn3163039). From the GTEx project (v8), we obtained gene-level count data in 13 brain regions, which will be made publicly available. We removed low-expressed genes (those genes for which fewer than three individuals had counts-per-million > 1) before normalization. We then normalized the RNA-seq raw counts using the trimmed mean of M-values normalization method^62^.

### Definitions of an individual-specific deconvolution model

The ultimate goal of the deconvolution is to estimate the cell type fraction *π*_*ik*_ of each cell type *k* in an individual *i*, treating the selected marker genes as data points. In each bulk sample *i*, we fit the NB model by regressing the bulk profile vector **y**_*i*_ on the estimated cell type profile matrix 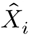, learned from snRNA-seq data.

We assume each gene-level quantification, *Y* (or *Y*_*gi*_ for a gene *g* on sample *i*), follows Negative Binomial (NB) distribution._^63^_Namely, we define the data likelihood of Y with the mean *μ* and over-dispersion *ϕ* parameters:

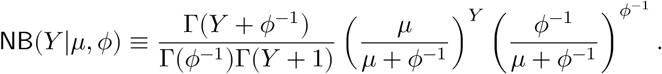

We define the NB model for the deconvolution problem:

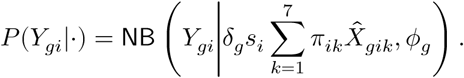

Here, we introduce auxiliary parameters, besides the *π* parameter:

- *s*_*i*_: sample-specific bias term for each individual *i* (easily estimable)
- *ϕ*_*g*_: over-dispersion parameter for each gene *g* (easily estimable)
- *δ*_*g*_: gene-specific bias term for each gene *g* in the bulk data

As for the first two parameters, we simply replace the sampling bias *s*_*i*_ with the sequencing depth of the bulk sample and find a suitable gene-level dispersion parameter *δ*_*g*_ using an empirical Bayes method implemented in `edgeR`^63^. However, finding a suitable *δ* value is non-trivial as this can be tightly dependent with *π* and is shared across all the samples. We discuss posterior inference algorithms in the next section.

### Reference cell-type models with the sample-specific covariates adjusted (steps 1 and 2)

From the snRNA-seq profiling followed by clustering analysis^6^, we construct a cell type-specific marker gene expression matrix *R*_*gik*_ (of gene *g*, sample *i*, cell type *k*), including the 171 marker genes (∼25 most differentially expressed in each cell type). Unlike conventional deconvolution methods^10,11^that directly use these marker gene profiles to estimate cell type fractions of bulk RNA-seq data, we adapt the cell type-specific marker gene model to the heterogeneity of biological and technical covariates.

We train each cell type *k*’s model by conducting the following NB regression model, *NB*(*R*_*gik*_|*l*_*i*_*μ*_*gik*_, *ψ*_*g*_) across *i* = 1, …, 48 individuals, where *l*_*i*_ is the library size of sample *i*. We further specify the mean function *μ*_*gik*_ as ln 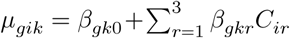 with the baseline activity *β*_*gk*0_ and the observed covariates *C*_*i*1_,*C*_2*i*_, *C*_*i*3_ correspond to the age, sex, and AD of an individual *i*, respectively.

We first estimate the overdispersion parameter *ψ*_*g*_ using `DESeq2`^64^. We then estimate the NB regression parameters using Stan^65^and construct the adjusted reference panel for a new sample *i* by plugging in the trained model parameters, 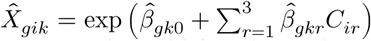. If all the coefficients (*β*) except the baseline were set to zero, our reference panel would be identical to the marker gene profiles used in the existing methods, but by including any non-zero effects of the known covariates, we prevent the marker genes from being influenced by these covariates in the subsequent deconvolution steps.

### Learning gene-specific bias between the bulk and snRNA-seq (step 3)

In our preliminary experiments, a brute-force parametric estimation method that directly estimate the posterior distribution of the bias and the cell-type fraction parameters often yielded poor results, e.g., high variance. Instead, we estimate *δ*_*g*_, assuming individual-level cell type fractions *π*_*ik*_ can be summarized average 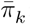:

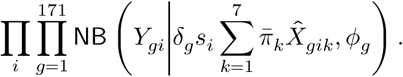

To estimate the average 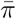, we leverage the subset of 69 “control” genes (Supplementary Fig. 1b, 1d) whose relative expression levels are robustly stable between the bulk RNA-seq and snRNA-seq data, and less variable across individuals:

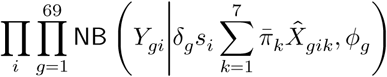

We optimize them in an EM algorithm by alternating between the two models: one for *δ* holding 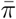 fixed and the other for 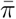 fixing *δ* values.

### Deconvolution to estimate the individual-specific cell type composition (step 4)

Provided that we have estimated auxiliary variables (*δ, s*, and *ϕ*) along with the parameters in the individual-specific reference cell type models (*β*), we resolve the individual-level cell type compositions (*π*_i*k*_) in Bayesian inference using using Stan^65^.

### Additional calibration step to compute the cell-level fraction estimates of cell types

To convert the transcript-level cell type fraction estimates to the composition of actual cell counts, we need to adjust a differential level of transcript abundance per cell across different cell types. Using the average number of transcripts 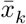 per cell within each cell type *k* in the snRNA-seq data, we reverse-engineer the cell-level fraction (*π*′) of each type that could have generated the estimate transcript-level fractions (*π*). We solve the following optimization for *π*′:

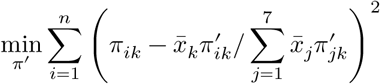

subject to 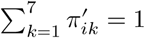 for each *i* and 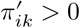.

### Pathway enrichment

We measure the impact of cell-type fractions on downstream transcript levels at the pathway-level. Within each pathway, and for each cell type, we compute gene-level z-scores that estimate significance of covariance between the cell-type fraction and the genes in the pathway. Say that we construct a test statistic for a pathway with m genes on a cell type *k*: we first standardize the cell-type fraction scores *p*_*ik*_ (for individualn *i* = 1, …, n, and cell type *k*) and gene expression *x*_*ig*_ (for an individual *i* and a gene *g*), and construct a gene-level score 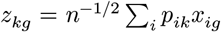. Combining these, we have test statistic 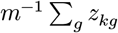 across *m* genes within each pathway. We estimate the null distribution by sample permutation along the individual axis.

### Genotype data imputation

We collected genotypes of 672,266 SNPs in 1,709 individuals from the Religious Orders Study (ROS) and the Memory and Aging Project (MAP)^46^in the GWAS for detecting cfGWAS hits. We mapped hg18 coordinates of SNPs (Affymetrix GeneChip 6.0) to hg19 coordinates, matching strands using publicly available information (http://www.well.ox.ac.uk/~wrayner/strand/GenomeWideSNP_6.na32-b37.strand.zip). We retained only those SNPs with MAF>0.05 and Hardy-Weinberg equilibrium (HWE) p-value>1e-04, computed based on 432 individuals who had all phenotype, genotype, and gene expression data. We imputed the genotypes by pre-phasing haplotypes based on the 1000 genome project^66^(phase I version 3) using SHAPEIT^67^. We then imputed SNPs in 5MB windows using IMPUTE2^68^with 100 Markov Chain Monte Carlo iterations and 10 burn-in iterations and retained only SNPs with MAF> 0.05 and imputation quality score>0.6. For the Mayo RNA-Seq project, we used a genotype dataset imputed by the Michigan Imputation Server^69^with the Haplotype Reference Consortium (hrc.r1.1.2016) panel^70^. The following documents provide more details about the Mayo dataset: https://www.synapse.org/#!Synapse:syn8650955.

### Polygenic risk scores

We modeled the polygenic risk *ρ*_*i*_ of an individual *i* as a weighted average of scaled genotype information^71^: *ρ*_*i*_ = ∑_*j*_ *G*_*ij*_*θ*_*j*_ where we take weighted average of genotype information *G*_*ij*_ (of individual *i* and SNP *j*) with the coefficients *θ*_*j*_ transferred from GWAS summary statistics data with the p-value threshold (p < 0.01) but the LD (linkage disequilibrium) structures decorrelated. Lacking individual-level phenotypes on all the available GWAS statistics, we fixed the p-value cutoff and the LD pruning steps were replaced with the decorrelation steps.^37,38^However, fine-tuning these parameters will only improve the performance.

### Sparse Bayesian regression to deconvolve tissue-level genetic effects into cell-type-specific eQTLs

We designed the deconvolved eQTL (deQTL) model from the following Bayesian generative scheme:

1. For each genetic variant *j* and cell type *k*, we sample unique multivariate eQTL effect sizes *θ*_*jk*_ *∼* a spike-slab prior.^72^(2) Each cell type *k* generates expression variation across individuals by a linear model *η*_*ik*_ = ∑_*j*_ *G*_*ij*_*θ*_*jk*_ on genetic information *G*_*ij*_ of each individual *i* in SNP *j*. (3) However, we only observe bulk gene expression profile 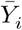 that is a mixture of cell type-specific genetic effects *η*_*ik*_ across the seven cell types, with some mixing proportion *π*_*k*_ (Fig. 6a). Provided that the estimate cell-type composition *π* is unbiased, we can model the mean of bulk profile as:

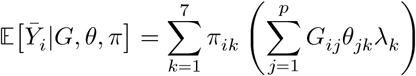

where we additionally include probabilistic loading factor *λ*_*k*_ ∈ (0, 1).

If we estimate the deQTL model SNP by SNP and cell-type by cell-type (*p* = 1), this model simply resorts to an interaction eQTL model^73^, testing non-zero-ness of the coefficient *θ*_*k*_ in Y_*i*_ *∼ θ*_*jk*_*π*_*ik*_ × *G*_*ij*_ without two singleton terms, which are *π* and *G*. In our multivariate model, we could include these non-interacting terms, but we only found such an over-parameterization was not as powerful as one might have expected. We concluded that these extra terms are rather unnecessary because these are likely to mediate the effect of cell-type-specific genetic variables by construction. It is widely accepted that effect size estimation of a causal path, while conditioning on an intermediate variable, can easily produce a biased result.^74^

### Simulation of bulk eQTL data using actual cell type composition and genotype matrix

We first select a causal cell type out of seven brain cell types where there are genetic effects on causal SNPs. In the Fig. 6b, we only show the results on the data simulated with three causal SNPs, but we varied the number of causal SNPs from 1 to 3. Our simulator generates gene expression data using the actual genotype matrix (*G*, standardized) and the deconvolved cell type estimations (*π*). We evaluated statistical power under the different level of total expression heritability (*h*^2^), varying from 5% to 40%. Provided that there is one cell type (out of total K=7) genetically-regulated with three causal SNPs, our simulator generates convolved gene expression profiles in the following steps.

1. For each celltype *k* ∈ [K] (K=7), if *k* is causal: we select three causal SNPs (*j*’s) uniformly at random and sample each genetic effect size θ _*jk*_ .𝒩 (0, (*K* / 3)^2^) For non-causal SNPs, we simply let the effect size *θ*_*jk*_ = 0. The deconvolved expression vector is constructed by a linear combination of the selected SNPs: **y**_*k*_ *∼ Gθ*_*k*_*π*_*k*_.
2. For the rest of non-causal cell type *l* ≠ *k*, we assign the expression vector **y**_*l*_ tonon-genetic signals by sampling from isotropic standard Gaussian distribution, and combine them by taking a weighted linear combination, **y**_0_ = ∑_*l*∈non-causal_ **y**_*l*_*π*_*l*_ except for the genetically regulated cell types.
3. We rescale **y**_0_ by multiplying a scaling factor to to achieve 𝕍 [***y*** _0_] = 𝕍 [*η* _*g*_] (1/ *h*^2^ − 1) to ensure that the simulated heritability to match with the assumed level, namely, *h*^2^ = 𝕍 [*η* _*g*_]/ (𝕍 [***y*** _0_] + 𝕍 [*η* _*g*_]).
4. The bulk RNA-seq data can be just a linear combination of these simulated celltype profiles: 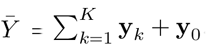.

### Competing deconvolved eQTL methods

We include comparison with other commonly used interaction QTL methods (Fig. 6):

- `deQTL` (this work): We fit multivariate deQTL model with stochastic variational Bayes inference algorithm. We then prioritize cell types in descending order of maximal posterior inclusion probability of genetic effects max 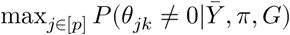, for each cell type *k*.
- `deQTL` (this work with additional terms): We prioritize cell types by the same procedure as above (deQTL) except that we added extra (and unnecessary) non-interaction terms of genotypes and cell types.
- Interaction QTL: We estimate the full set of p-values for conventional interaction QTL analysis using `lm(y ∼ cell type * genotype + cell type + genotype)` in `R`. We then summarize each cell type’s score by minimum p-values across SNPs within each cell type. Prioritize the cell types in the ascending order of the minimal p-values.

### Immunostaining validation of the predicted cell-type fractions

Fixed human brain tissue (prefrontal cortex, BA10) was sectioned at 50 m using a vibratome (Leica). The sections were boiled in IHC Antigen Retrieval Solution (ThermoFisher Scientific; catalog number 00-4955-58) containing 0.05% Tween-20 for 10 minutes and then placed in PBS for 20 minutes at room temperature. After washing with ddH2O (three times 15 minutes) followed by one wash with PBS for 15 minutes, the brain sections were incubated in quenching solution (50mM ammonium acetate, 100mM CuSO4) at room temperature overnight. After washing with ddH20 (one wash for 15 minutes) and PBS (three times 15 minutes), the sections were permeabilized in PBS containing 0.3% Triton X-100 for 10 minutes and blocked in PBS containing 0.3% Triton X-100 and 5% normal donkey serum at room temperature for 2 h. The sections were incubated for 2 hours at room temperature in primary antibody in PBS with 0.3% Triton X-100 and 5% normal donkey serum. Primary antibodies were an anti-GFAP antibody (1:100; Abcam; ab53554, Goat polyclonal) and anti-Iba1 Antibody (1:500; Synaptic Systems; Cat. No. 234 004, Polyclonal Guinea pig antiserum). The sections were washed with PBS containing 0.3% Triton X-100 and 5% normal donkey serum at room temperature (four times 15 minutes) and then incubated with secondary antibodies (dilution 1:2000) for 2 hours at room temperature. Primary antibodies were visualized with Alexa-Fluor 488 and Alexa-Fluor 594 antibodies (Molecular Probes), and cell nuclei visualized with Hoechst 33342 (Sigma-Aldrich; 94403). The sections were washed with PBS containing 0.3% Triton X-100 and 5% normal donkey serum at room temperature (four times 15 minutes) and then mounted on Fisherbrand (TM) Superfrost (TM) Plus Microscope Slides in ProLong (TM) Gold Antifade Mountant. Images were acquired using a confocal microscope (LSM 710; Zeiss) with a 20x or 40x objective, and cell numbers were quantified using Imaris 8.3.1.

**Table 1.**
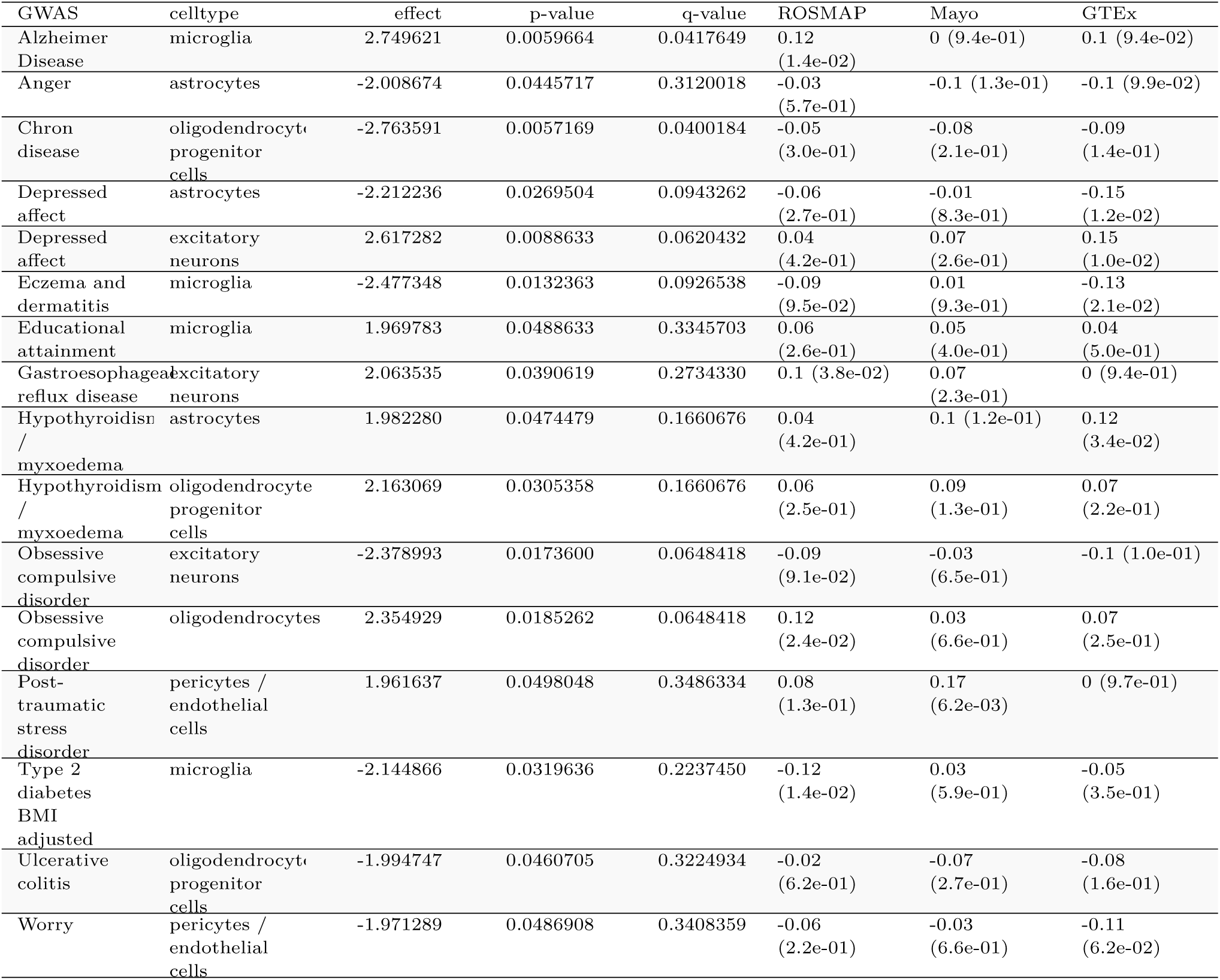
Polygenic risk scores significantly correlated with the estimated cell type fractions (p-value < 0.05).

## Supplementary Figures

**Supplementary Figure 1.**
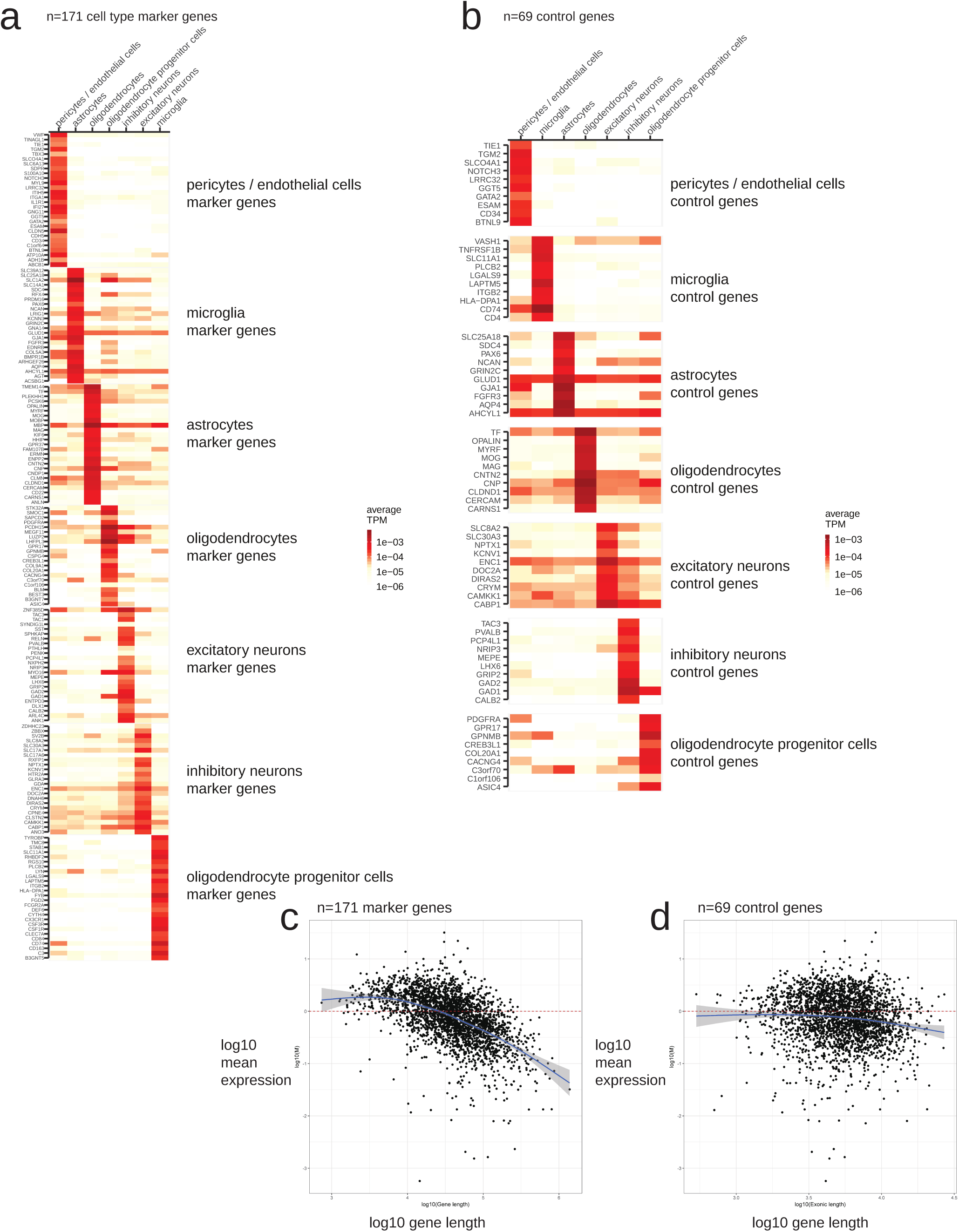
Average expression profiles of cell type marker genes in snRNA-seq data. (a) A total set of marker genes’ average expression profiles in snRNA-seq data. (b) The average expression profiles of the control genes in snRNA-seq data. (c) There exists gene-length bias in the total set of marker genes. (d) We select the control genes to avoid the gene length bias.

**Supplementary Figure 2.**
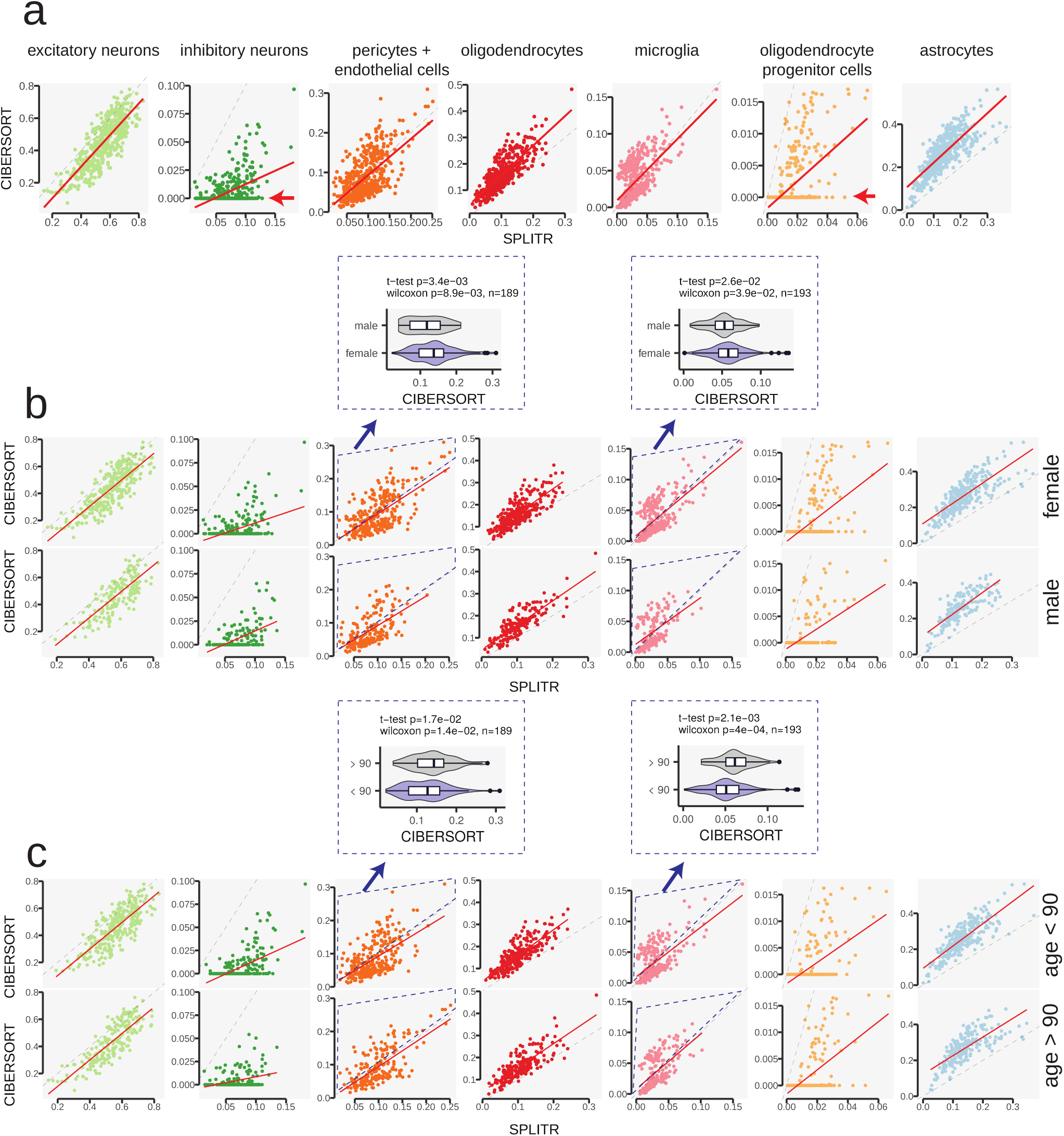
Comparison with CIBERSORT. (a) We compared the performance of our cell-type deconvolution methods (SPLITR) with the other method (CIBERSORT^10^), providing the same maker gene profile matrix. (b) We highlight the difference between two methods in pericytes and microglia can be explained by sex (male vs. female) difference in the bulk data. (c) We highlight the difference between the two methods in pericytes and microglia can be explained by age difference in the bulk data.

**Supplementary Figure 3.**
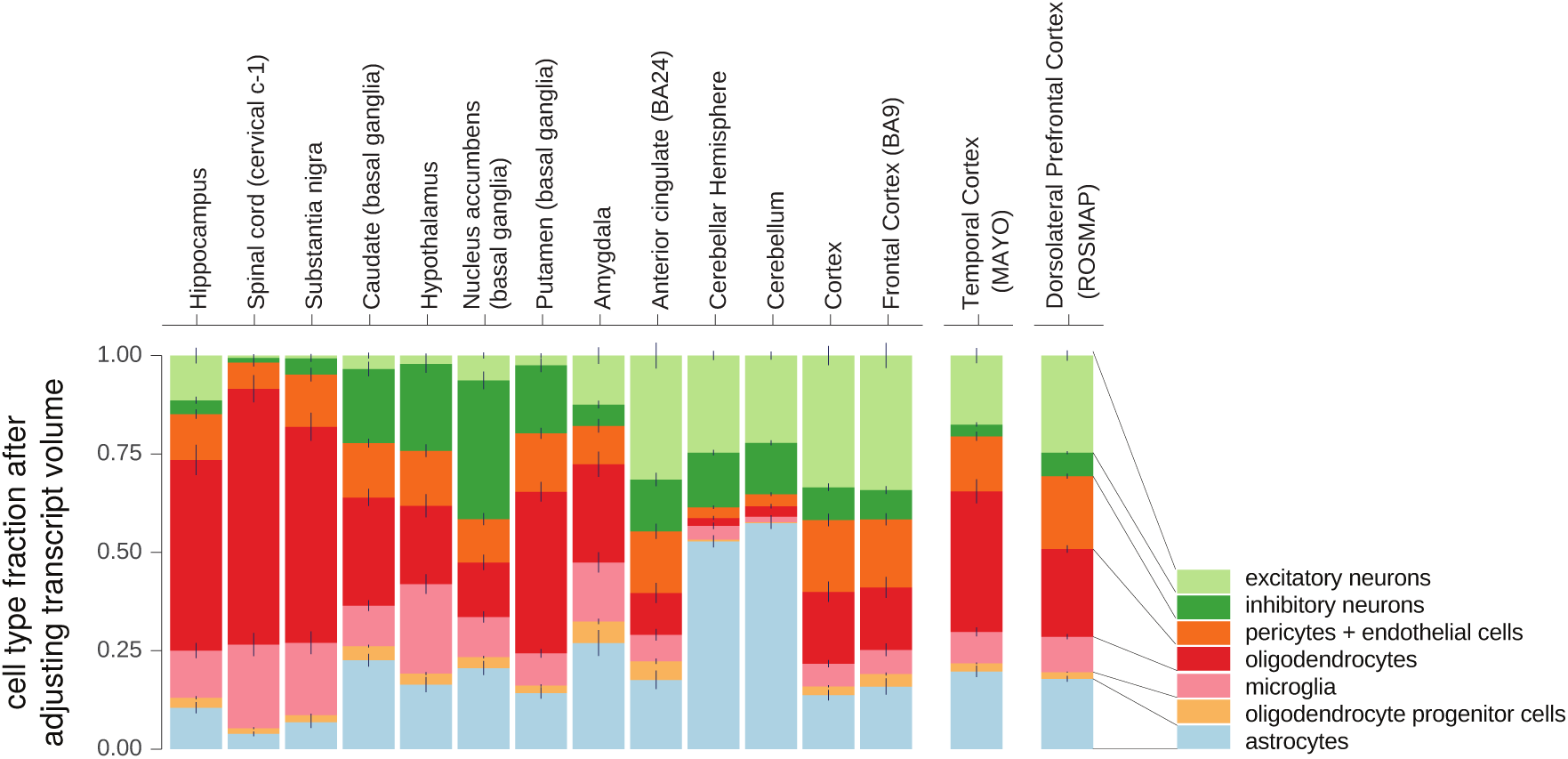
Average cell type fractions estimated at cell-count-level.

**Supplementary Figure 4.**
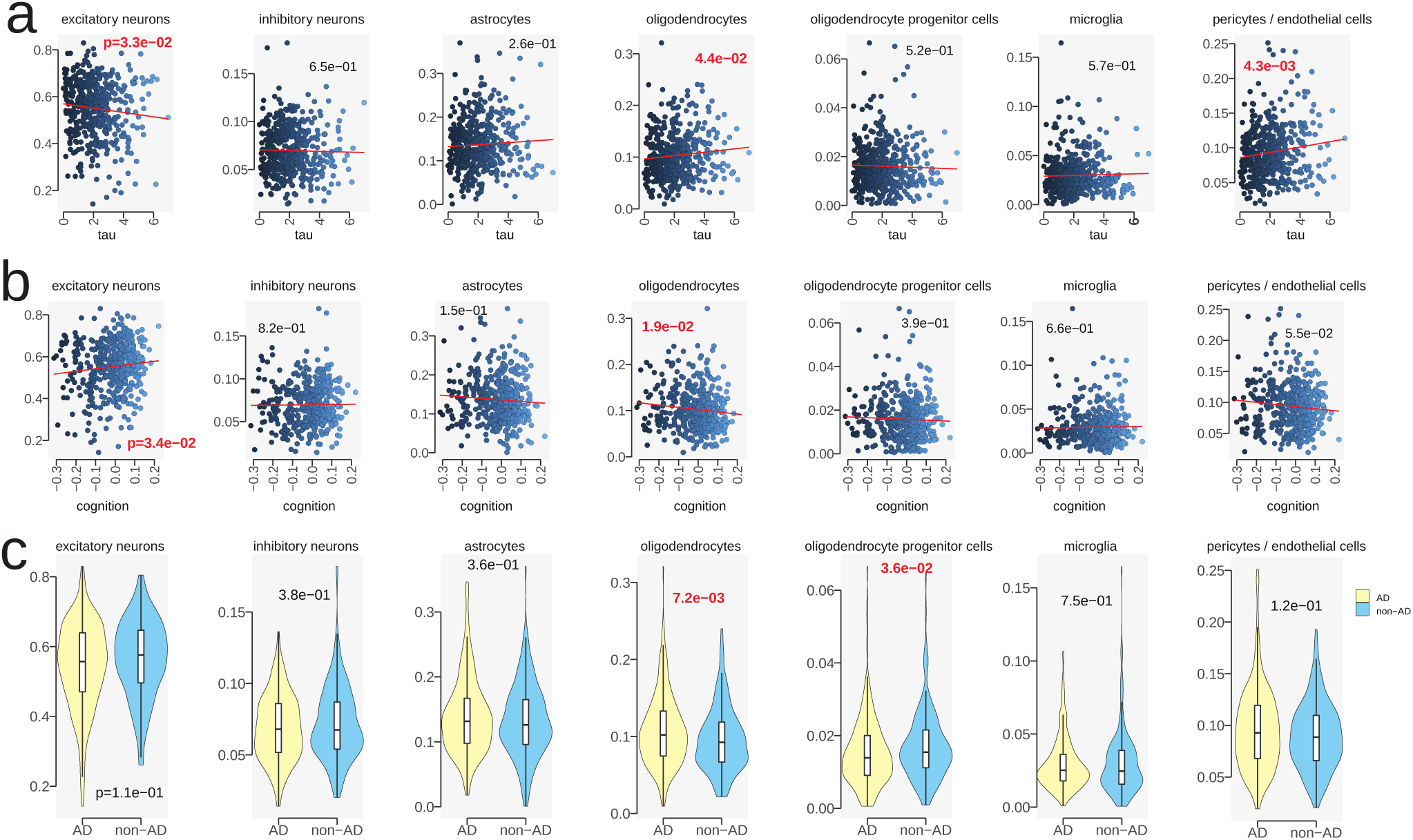
Correlation between cell-type fractions in the brain samples with known pathological variables. Y-axis: cell type composition; X-axis: pathological variables. (a) X-axis: amyloid-beta (in square root). (b) X-axis: neurofibrillary tangles tau protein (in square root). (c) Cell type compositions change between AD and non-AD samples.

**Supplementary Figure 5.**
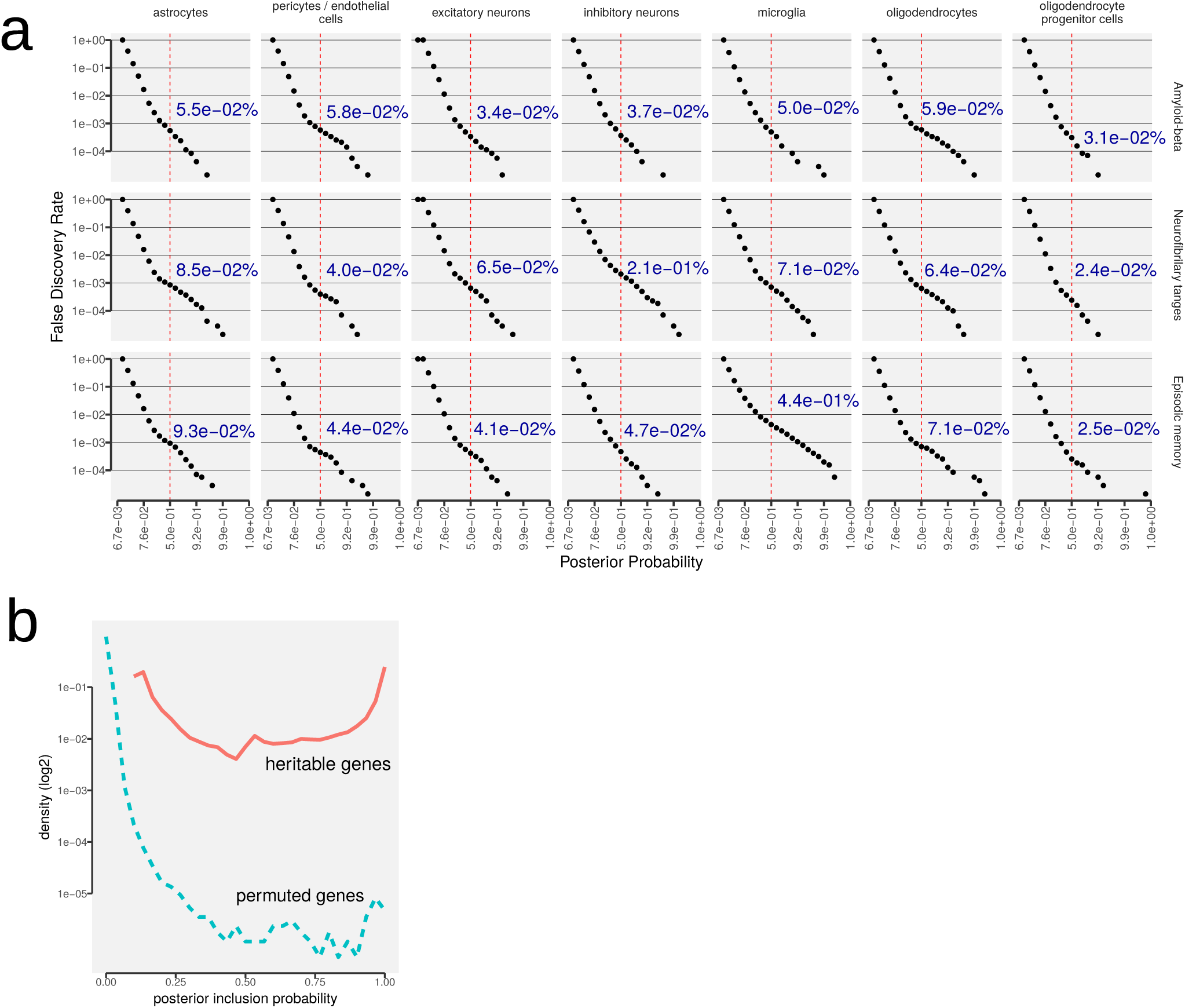
Empirically-calibrated false discovery rates of non-zero effects. (a) For the cell-type-specific gene models, we constructed the null data by Freedman-Lane permutation^39^. X-asis: posterior probability cutoff; y-axis: false discovery rate. (b) For the deQTL models, we constructed the null data by permuting samples after adjusting non-genetic factors. X-asis: posterior probability cutoff; y-axis: density (log2 scaled).

**Supplementary Figure 6.**
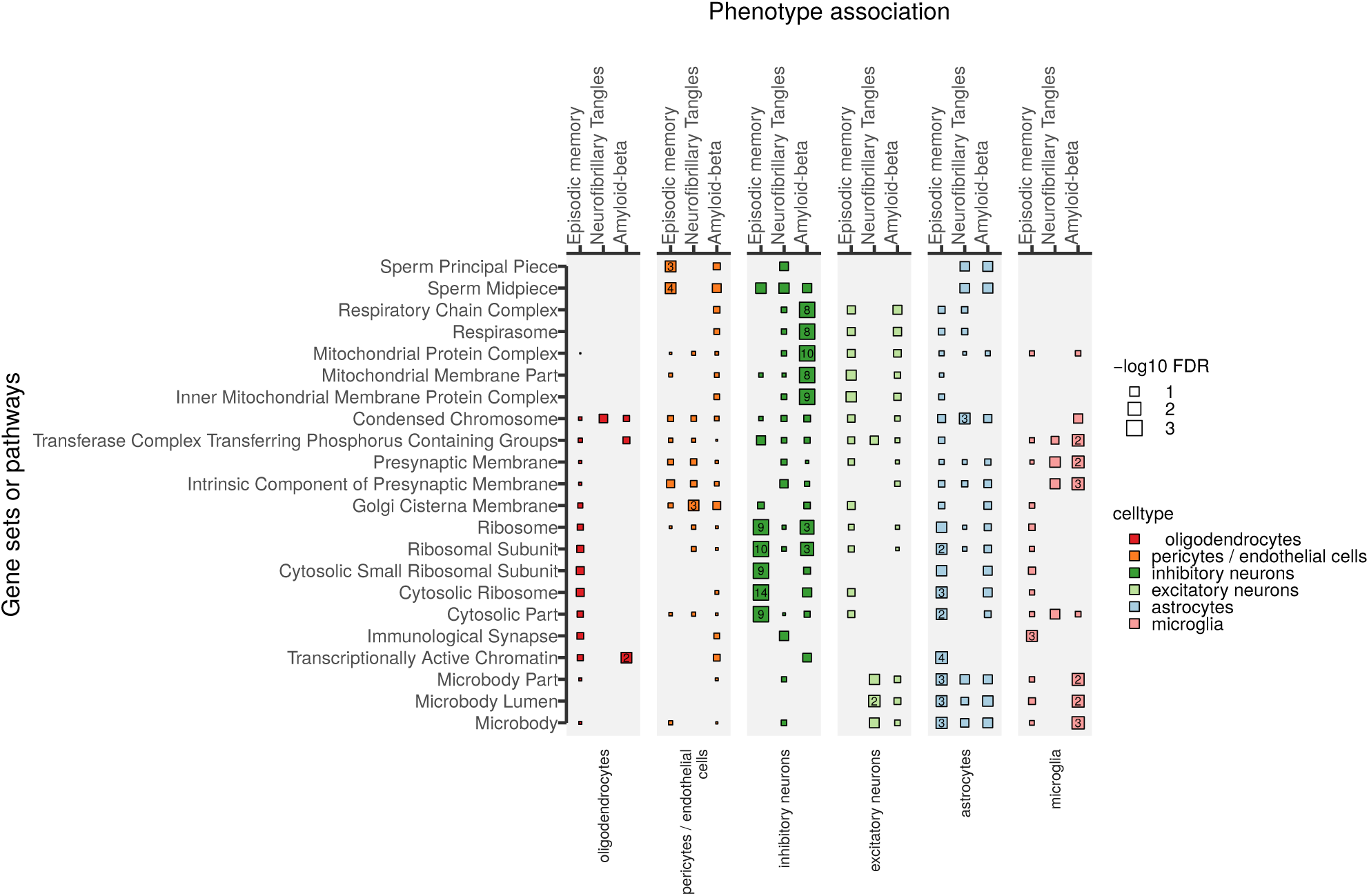
Cell-type-specific gene ontology (cellular component) enrichment results for the significantly-associated genes. X-asis: cell types; y-axis: keywords (gene sets).

**Supplementary Figure 7.**
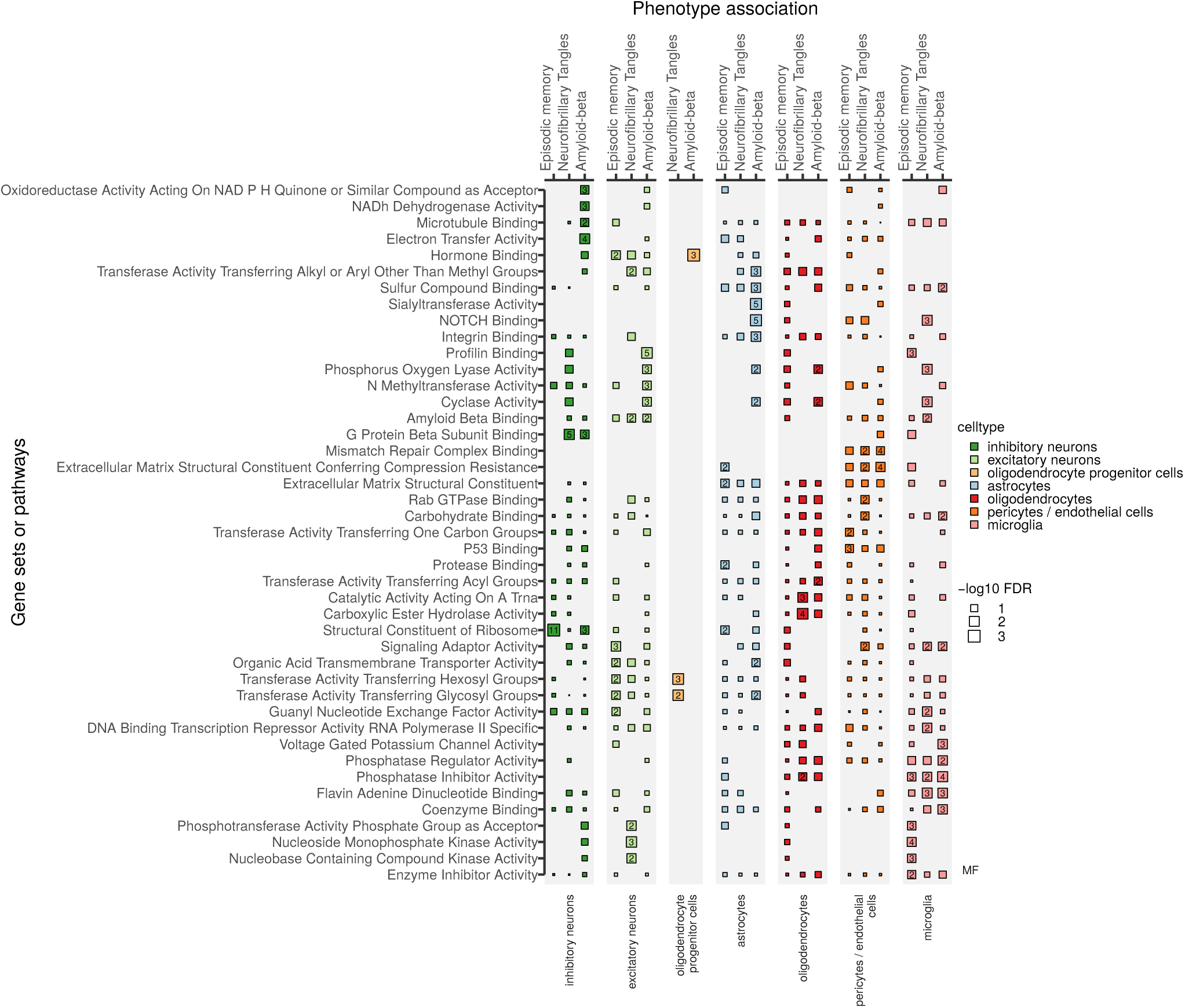
Cell-type-specific gene ontology (molecular function) enrichment results for the significantly-associated genes. X-asis: cell types; y-axis: keywords (gene sets).

## Supplementary Tables

**Supplementary Table 1.**
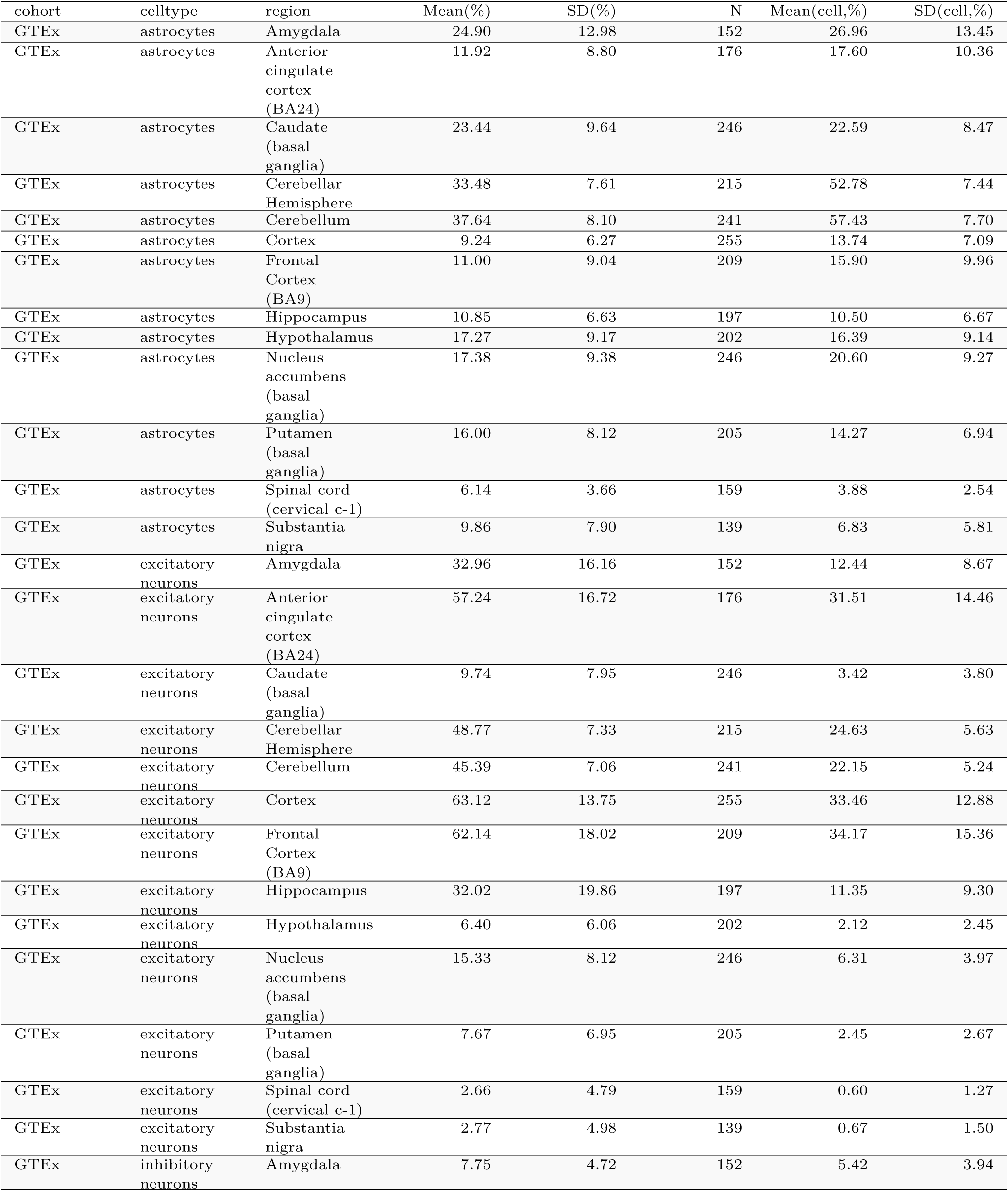
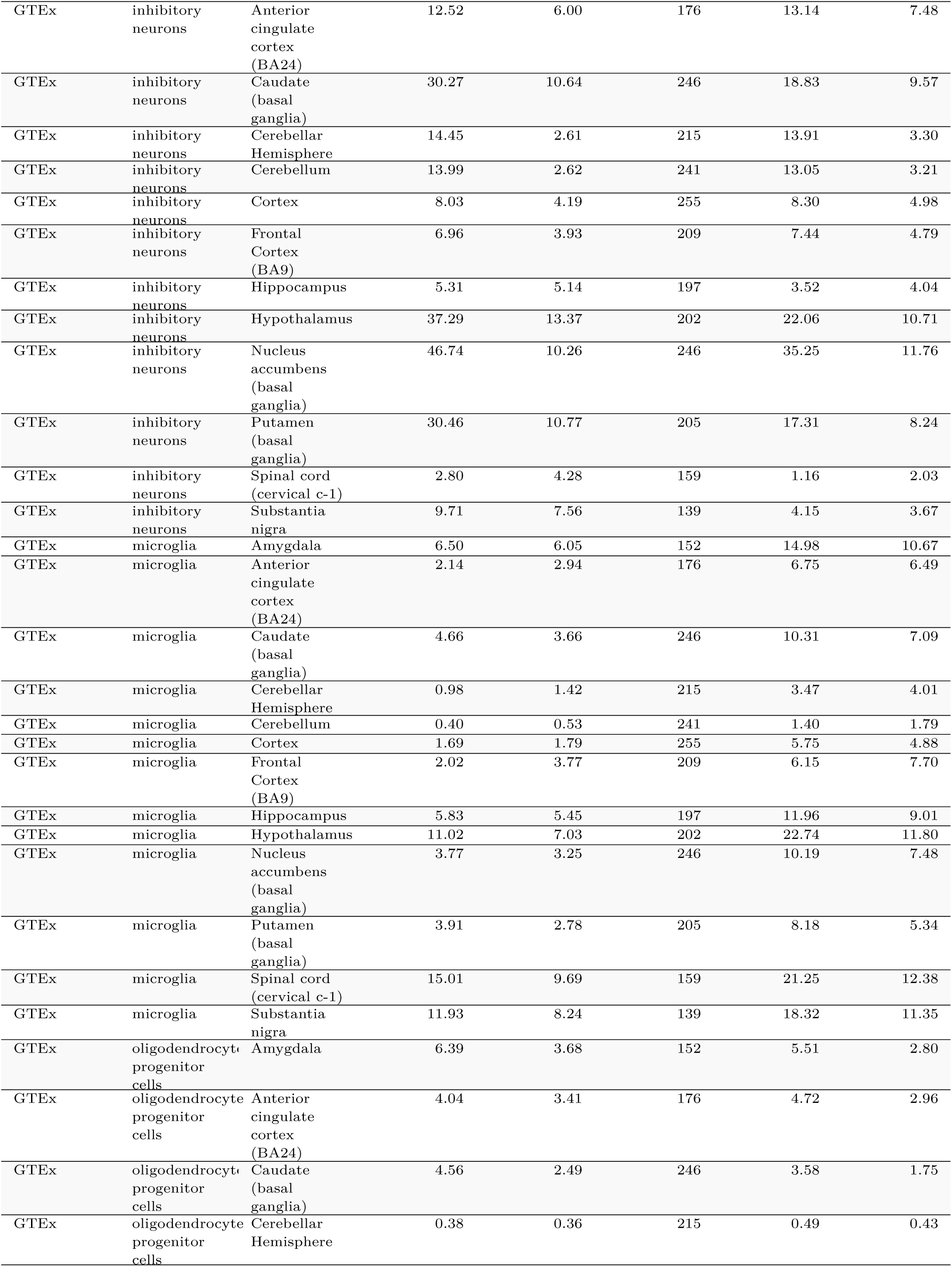
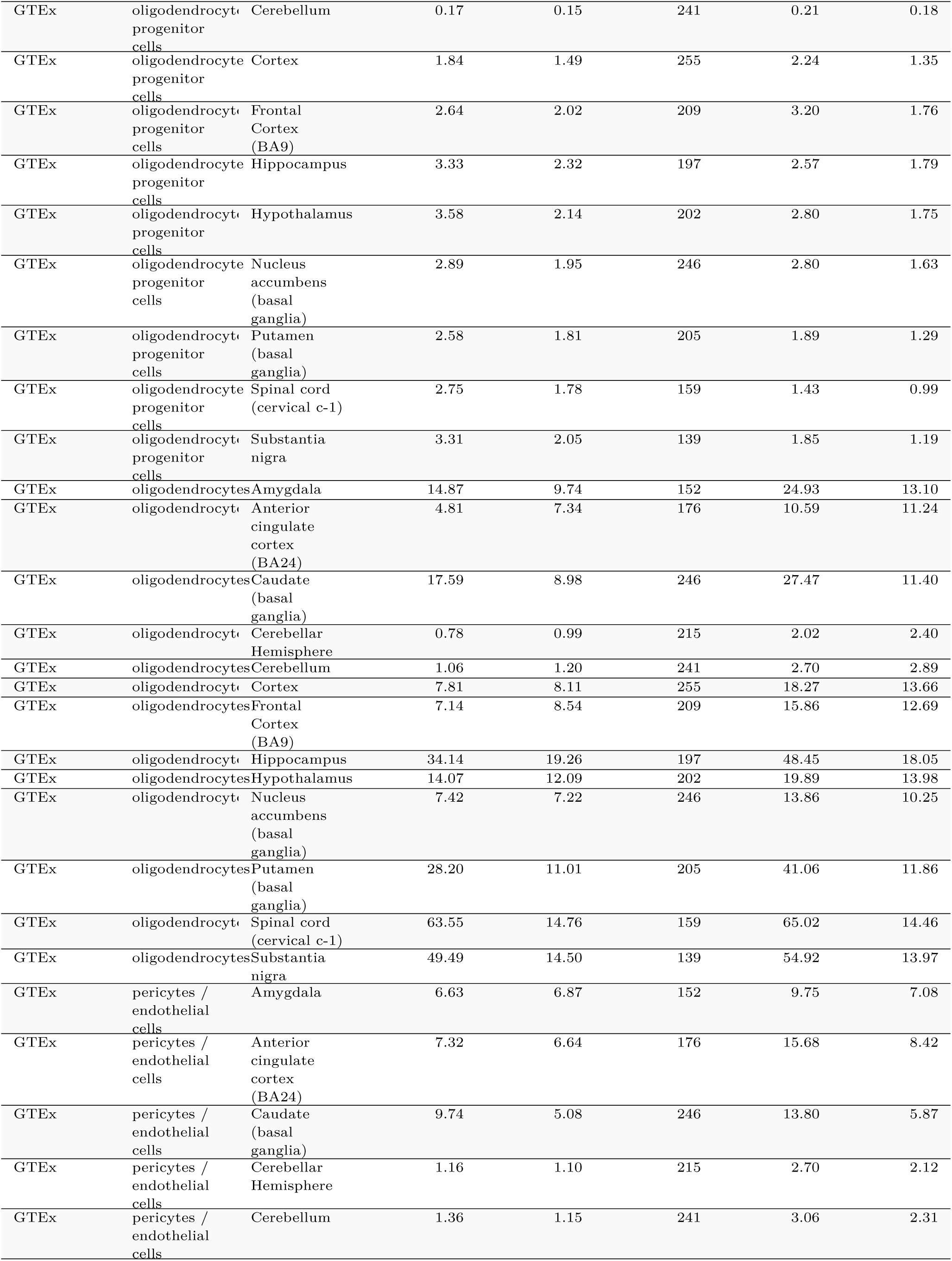
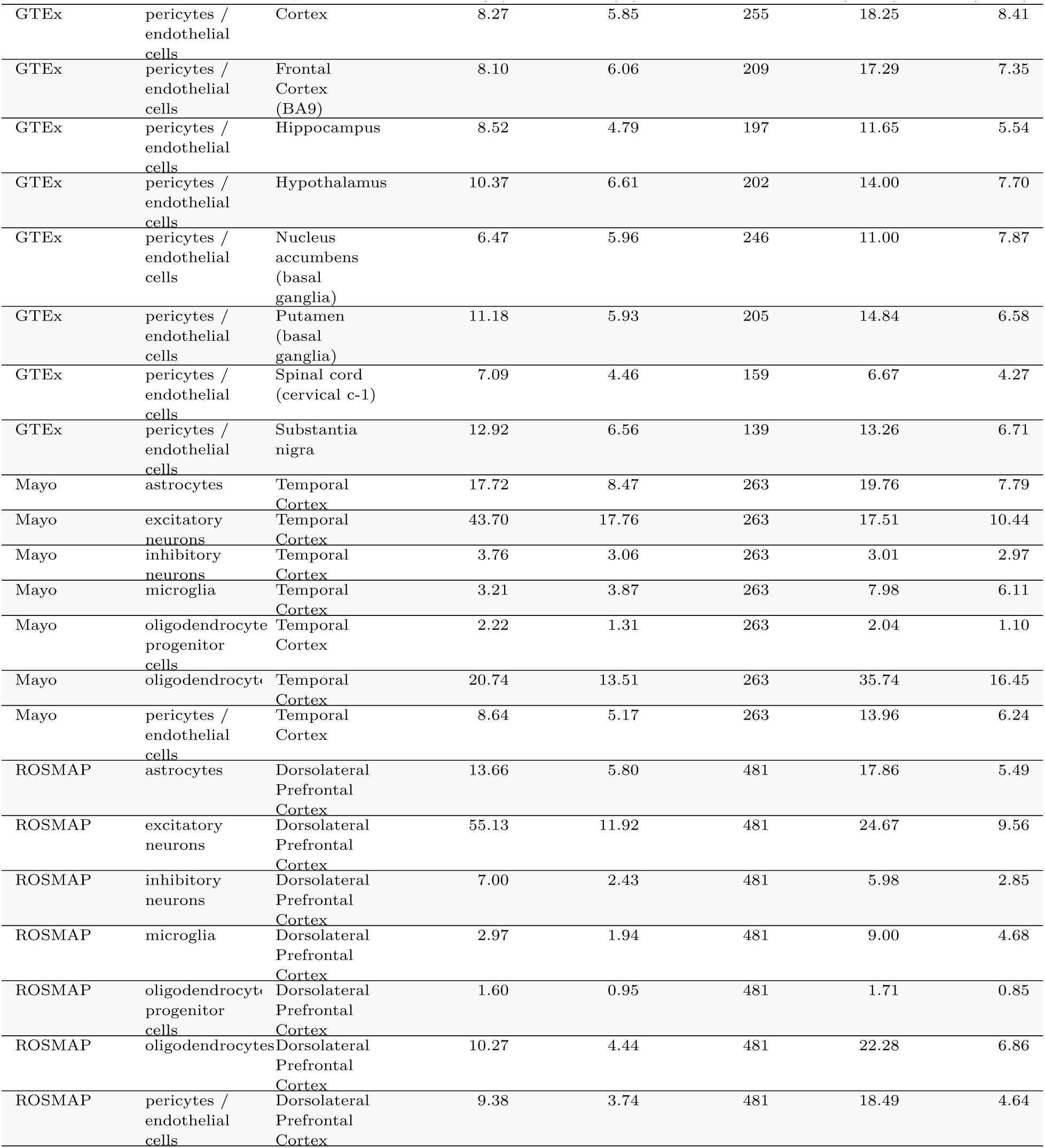
Summary of cell type deconvolution results

**Supplementary Table 2.** 56 GWAS traits analyzed in this work

| GWAS.traits | Source | N | Ncase | Ncontrol |
| --- | --- | --- | --- | --- |
| Alzheimer Disease | Jansen (2018) | 455,258 | 71,880 | 383,378 |
| Anger | Watanabe (2018) | NA | NA | 57,010 |
| Annoyance | Watanabe (2018) | 126,198 | NA | NA |
| asthma | Watanabe (2018) | 385,822 | 44,301 | 341,521 |
| Atopic dermatitis | Paternoster (2011) | 26,171 | 5,606 | 20,565 |
| Attention-deficit/hyperactivity disorder | Demontis (2017) | 53,293 | 19,099 | 34,194 |
| Autism spectrum disorder | Weiner (2017) | 46,350 | 18,381 | 27,969 |
| Automobile speeding propensity | Karlsson Linner (2019) | 404,291 | NA | NA |
| Basal metabolic rate | Watanabe (2018) | 379,821 | NA | NA |
| Bipolar and depression | Watanabe (2018) | 93,296 | 25,873 | 67,423 |
| Bipolar disorder | Stahl (2017) | 74,194 | 20,129 | 54,065 |
| Body fat percentage | Watanabe (2018) | 379,615 | NA | NA |
| Body mass index | Yengo (2018) | 700,000 | NA | NA |
| Cholesterol medication | Watanabe (2018) | 176,050 | 40,433 | 135,617 |
| Chron disease | Franke (2010) | 21,389 | 6,333 | 15,056 |
| Cognitive performance | Lee (2018) | 257,828 | NA | NA |
| Depressed affect | Nagel (2018) | 357,957 | NA | NA |
| Depression | Watanabe (2018) | 289,307 | 22,055 | 267,252 |
| Diabetes | Watanabe (2018) | 385,420 | 18,483 | 366,937 |
| Drinks per week | Karlsson Linner (2019) | 414,343 | NA | NA |
| Eczema and dermatitis | Watanabe (2018) | 9,381 | 279,476 | 289,307 |
| Educational attainment | Lee (2018) | 766,345 | NA | NA |
| Ever smoker | Karlsson Linner (2019) | 518,633 | NA | NA |
| Extreme irritability | Watanabe (2018) | 32,229 | 90,662 | 122,891 |
| Gallstones | Watanabe (2018) | 300,791 | 11,632 | 289,159 |
| Gastritis and duodenitis | Watanabe (2018) | 300,791 | 14,477 | 286,314 |
| Gastroesophageal reflux disease | Watanabe (2018) | 300,791 | 12,011 | 288,780 |
| Hayfever / allergic rhinitis | Watanabe (2018) | 289,307 | 22,057 | 267,250 |
| Hayfever / allergic rhinitis / eczema | Watanabe (2018) | 385,822 | 89,380 | 296,442 |
| High blood pressure | Watanabe (2018) | 385,699 | 103,381 | 282,318 |
| High cholesterol | Watanabe (2018) | 289,307 | 46,932 | 242,375 |
| Hypothyroidism / myxoedema | Watanabe (2018) | 289,307 | 18,740 | 270,567 |
| Inflammatory bowel disease | Liu (2015) | 96,486 | 9,846 | 86,640 |
| Insomnia | Jansen (2019) | 386,533 | 109,402 | 277,131 |
| Irritability | Watanabe (2018) | 369,232 | 104,545 | 264,687 |
| Irritable argument | Watanabe (2018) | 22,783 | 103,513 | 126,296 |
| Major depressive disorder | Watanabe (2018) | 244,890 | 10,402 | 234,488 |
| Major depressive disorder | Ripke (2013) | 173,005 | 59,851 | 113,154 |
| Number of sexual partners | Karlsson Linner (2019) | 370,711 | NA | NA |
| Obsessive compulsive disorder | Stewart (2013) and Mattheisen (2015) | 9,725 | 2,688 | 7,037 |
| Osteoarthritis (meta) | Zengini (2018) | 327,918 | 30,727 | 297,191 |
| Osteoarthritis (UKB) | Watanabe (2018) | 289,307 | 31,545 | 257,762 |
| Osteoarthritis of hip | Watanabe (2018) | 300,791 | 9,873 | 290,918 |
| Osteoarthritis of knee | Watanabe (2018) | 300,791 | 12,508 | 288,283 |
| Post-traumatic stress disorder | Duncan (2017) | 9,537 | 2,424 | 7,113 |
| Rheumatoid arthritis | Okaba (2014) | 103,638 | 29,880 | 73,758 |
| Risk tolerance | Karlsson Linner (2019) | 466,571 | NA | NA |
| Schizophrenia | Ripke (2014) | 87,491 | 33,426 | 54,065 |
| Systemic lupus erythematosus | Bentham (2015) | 23,210 | 7,219 | 15,991 |
| The 1st PC of the four<br>risky behaviors | Karlsson Linner (2019) | 315,894 | NA | NA |
| Traumatic depression | Watanabe (2018) | 71,568 | 52,198 | 19,370 |
| Type 2 diabetes (meta) | Mahajan (2018) | 898,130 | 74,124 | 824,006 |
| Type 2 diabetes (UKB) | Watanabe (2018) | 244,890 | 16,673 | 228,217 |
| Type 2 diabetes BMI<br>adjusted | Mahajan (2018) | 898,130 | 74,124 | 824,006 |
| Ulcerative colitis | Anderson (2011) | 26,405 | 6,687 | 19,718 |
| Worry | Nagel (2018) | 348,219 | NA | NA |

**Supplementary Table 3.** Number of tissue-level eQTL and cell-type-specific deQTL SNPs and genes.

| cell type | #eQTLs | #eQTLs<br>cell-type<br>only | #eQTLs<br>cell-type<br>only (%) | #genes | #genes<br>cell-type<br>only | #genes<br>cell-type<br>only (%) | color |
| --- | --- | --- | --- | --- | --- | --- | --- |
| brain tissue | 7,783 | 0 | 0 | 5,586 | 0 | 0 | #DDDDDD |
| cell-type<br>combined | 4,757 | 1,252 | 26 | 3,869 | 1,182 | 31 | #FFCC00 |
| inhibitory<br>neurons | 1,966 | 410 | 21 | 1,727 | 391 | 23 | #389D34 |
| excitatory<br>neurons | 1,259 | 239 | 19 | 1,071 | 230 | 21 | #BDE29C |
| oligodendrocyte | 710 | 228 | 32 | 692 | 229 | 33 | #E01F25 |
| astrocytes | 588 | 147 | 25 | 556 | 147 | 26 | #A8CEE1 |
| microglia | 364 | 176 | 48 | 357 | 173 | 48 | #F89C9B |
| pericytes /<br>endothelial<br>cells | 319 | 78 | 24 | 307 | 78 | 25 | #FD7C21 |
| oligodendrocyte<br>progenitor<br>cells | 24 | 11 | 46 | 23 | 11 | 48 | #FBBC74 |

## References

1. Lambert, J. C. et al. Meta-analysis of 74,046 individuals identifies 11 new susceptibility loci for Alzheimer’s disease. Nat. Genet. 45, 1452–1458 (2013).

2. Jansen, I. E. et al. Genome-wide meta-analysis identifies new loci and functional pathways influencing alzheimer’s disease risk. Nat. Genet. (2019).

3. Mostafavi, S. et al. A molecular network of the aging human brain provides insights into the pathology and cognitive decline of alzheimer’s disease. Nat. Neurosci. 21, 811–819 (2018).

4. De Jager, P. L. et al. Alzheimer’s disease: early alterations in brain DNA methylation at ANK1, BIN1, RHBDF2 and other loci. Nat. Neurosci. 17, 1156–1163 (2014).

5. Zhang, B. et al. Integrated systems approach identifies genetic nodes and networks in late-onset Alzheimer’s disease. Cell 153, 707–720 (2013).

6. Mathys, H. et al. Single-cell transcriptomic analysis of alzheimer’s disease. Nature 570, 332–337 (2019).

7. Lake, B. B. et al. Neuronal subtypes and diversity revealed by single-nucleus RNA sequencing of the human brain. Science 352, 1586–1590 (2016).

8. Ng, B. et al. An xQTL map integrates the genetic architecture of the human brain’s transcriptome and epigenome. Nat. Neurosci. 20, 1418–1426 (2017).

9. GTEx Consortium et al. Genetic effects on gene expression across human tissues. Nature 550, 204–213 (2017).

10. Newman, A. M. et al. Robust enumeration of cell subsets from tissue expression profiles. Nat. Methods 12, 453–457 (2015).

11. Aran, D., Hu, Z. & Butte, A. J. XCell: Digitally portraying the tissue cellular heterogeneity landscape. Genome Biol. 18, 220 (2017).

12. Schelker, M. et al. Estimation of immune cell content in tumour tissue using single-cell RNA-seq data. Nat. Commun. 8, 2032 (2017).

13. Tirosh, I. et al. Dissecting the multicellular ecosystem of metastatic melanoma by single-cell RNA-seq. Science 352, 189–196 (2016).

14. Trapnell, C. et al. The dynamics and regulators of cell fate decisions are revealed by pseudotemporal ordering of single cells. Nat. Biotechnol. 32, 381–386 (2014).

15. Frishberg, A. et al. Cell composition analysis of bulk genomics using single-cell data. Nat. Methods 16,n 327–332 (2019).

16. Zhang, Y. et al. Purification and characterization of progenitor and mature human astrocytes reveals transcriptional and functional differences with mouse. Neuron 89, 37–53 (2016).

17. Bennett, D. A. et al. The rush memory and aging project: Study design and baseline characteristics of the study cohort. Neuroepidemiology 25, 163–175 (2005).

18. Allen, M. et al. Human whole genome genotype and transcriptome data for alzheimer’s and other neurodegenerative diseases. Sci Data 3, 160089 (2016).

19. Bartheld, C. S. von, Bahney, J. & Herculano-Houzel, S. The search for true numbers of neurons and glial cells in the human brain: A review of 150 years of cell counting. J. Comp. Neurol. 524, 3865–3895 (2016).

20. Lyck, L. et al. An empirical analysis of the precision of estimating the numbers of neurons and glia in human neocortex using a fractionator-design with sub-sampling. J. Neurosci. Methods 182, 143–156 (2009).

21. Andrade-Moraes, C. H. et al. Cell number changes in alzheimer’s disease relate to dementia, not to plaques and tangles. Brain 136, 3738–3752 (2013).

22. Sahara, S., Yanagawa, Y., O’Leary, D. D. M. & Stevens, C. F. The fraction of cortical GABAergic neurons is constant from near the start of cortical neurogenesis to adulthood. J. Neurosci. 32, 4755–4761 (2012).

23. Chen-Plotkin, A. S. et al. TMEM106B, the risk gene for frontotemporal dementia, is regulated by the microRNA-132/212 cluster and affects progranulin pathways. J. Neurosci. 32, 11213–11227 (2012).

24. Van Deerlin, V. M. et al. Common variants at 7p21 are associated with frontotemporal lobar degeneration with TDP-43 inclusions. Nat. Genet. 42, 234–239 (2010).

25. Vass, R. et al. Risk genotypes at TMEM106B are associated with cognitive impairment in amyotrophic lateral sclerosis. Acta Neuropathol. 121, 373–380 (2011).

26. Zee, J. van der et al. TMEM106B is associated with frontotemporal lobar degeneration in a clinically diagnosed patient cohort. Brain 134, 808–815 (2011).

27. Rhinn, H. & Abeliovich, A. Differential aging analysis in human cerebral cortex identifies variants in TMEM106B and GRN that regulate aging phenotypes. Cell Syst 4, 404–415.e5 (2017).

28. Baker, M. et al. Mutations in progranulin cause tau-negative frontotemporal dementia linked to chromosome 17. Nature 442, 916–919 (2006).

29. Zhou, X., Sun, L., Brady, O. A., Murphy, K. A. & Hu, F. Elevated TMEM106B levels exaggerate lipofuscin accumulation and lysosomal dysfunction in aged mice with progranulin deficiency. Acta Neuropathol Commun 5, 9 (2017).

30. Gallagher, M. D. et al. A Dementia-Associated risk variant near TMEM106B alters chromatin architecture and gene expression. Am. J. Hum. Genet. 101, 643–663 (2017).

31. Satoh, J.-I. et al. TMEM106B expression is reduced in alzheimer’s disease brains. Alzheimers. Res. Ther. 6, 17 (2014).

32. Busch, J. I. et al. Increased expression of the frontotemporal dementia risk factor TMEM106B causes c9orf72-dependent alterations in lysosomes. Hum. Mol. Genet. 25, 2681–2697 (2016).

33. Vullhorst, D. et al. Selective expression of ErbB4 in interneurons, but not pyramidal cells, of the rodent hippocampus. J. Neurosci. 29, 12255–12264 (2009).

34. Wen, L. et al. Neuregulin 1 regulates pyramidal neuron activity via ErbB4 in parvalbumin-positive interneurons. Proc. Natl. Acad. Sci. U. S. A. 107, 1211–1216 (2010).

35. Chu, C.-S. et al. The DAOA gene is associated with schizophrenia in the taiwanese population. Psychiatry Res. 252, 201–207 (2017).

36. Ozburn, A. R. et al. NPAS2 regulation of Anxiety-Like behavior and GABAA receptors. Front. Mol. Neurosci. 10, 360 (2017).

37. Baker, E. et al. POLARIS: Polygenic LD-adjusted risk score approach for set-based analysis of GWAS data. Genet. Epidemiol. 42, 366–377 (2018).

38. Chun, S. et al. Non-parametric polygenic risk prediction using partitioned GWAS summary statistics. bioRxiv (2019).

39. Freedman, D. & Lane, D. A Nonstochastic Interpretation of Reported Significance Levels. J. Bus. Econ. Stat. 1, 292–298 (1983).

40. Liberzon, A. et al. Molecular signatures database (MSigDB) 3.0. Bioinformatics 27, 1739–1740 (2011).

41. Jaramillo-Merchán, J. et al. Mesenchymal stromal-cell transplants induce oligodendrocyte progenitor migration and remyelination in a chronic demyelination model. Cell Death Dis. 4, e779 (2013).

42. Rivera, F. J. et al. Aging restricts the ability of mesenchymal stem cells to promote the generation of oligodendrocytes during remyelination. Glia 67, 1510–1525 (2019).

43. Rajendran, L. & Paolicelli, R. C. Microglia-Mediated synapse loss in alzheimer’s disease. J. Neurosci. 38, 2911–2919 (2018).

44. González-Reyes, R. E., Nava-Mesa, M. O., Vargas-Sánchez, K., Ariza-Salamanca, D. & Mora-Muñoz, L. Involvement of astrocytes in alzheimer’s disease from a neuroinflammatory and oxidative stress perspective. Front. Mol. Neurosci. 10, 427 (2017).

45. Park, Y. et al. A bayesian approach to mediation analysis predicts 206 causal target genes in alzheimer’s disease. bioRxiv 219428 (2017).

46. De Jager, P. L. et al. A genome-wide scan for common variants affecting the rate of age-related cognitive decline. Neurobiol. Aging 33, 1017.e1–1017.e15 (2012).

47. Stewart, S. E. et al. Genome-wide association study of obsessive-compulsive disorder. Mol. Psychiatry 18, 788–798 (2013).

48. Mattheisen, M. et al. Genome-wide association study in obsessive-compulsive disorder: Results from the OCGAS. Mol. Psychiatry 20, 337–344 (2015).

49. Weiner, D. J. et al. Polygenic transmission disequilibrium confirms that common and rare variation act additively to create risk for autism spectrum disorders. Nat. Genet. 49, 978–985 (2017).

50. Zengini, E. et al. Genome-wide analyses using UK biobank data provide insights into the genetic architecture of osteoarthritis. Nat. Genet. 50, 549–558 (2018).

51. Lee, J. J. et al. Gene discovery and polygenic prediction from a genome-wide association study of educational attainment in 1.1 million individuals. Nat. Genet. 50, 1112–1121 (2018).

52. Karlsson Linnér, R. et al. Genome-wide association analyses of risk tolerance and risky behaviors in over 1 million individuals identify hundreds of loci and shared genetic influences. Nat. Genet. 51, 245–257 (2019).

53. Anderson, C. A. et al. Meta-analysis identifies 29 additional ulcerative colitis risk loci, increasing the number of confirmed associations to 47. Nat. Genet. 43, 246–252 (2011). view of pleiotropy and genetic architecture in complex traits. bioRxiv\ (2018).

54. Watanabe, K. et al. A global view of pleiotropy and genetic architecture in complex traits. bioRxiv (2018).

55. Zhang, B. et al. Integrated systems approach identifies genetic nodes and networks in late-onset alzheimer’s disease. Cell 153, 707–720 (2013).

56. Heppner, F. L., Ransohoff, R. M. & Becher, B. Immune attack: The role of inflammation in alzheimer disease. Nat. Rev. Neurosci. 16, 358–372 (2015).

57. Crotti, A. et al. BIN1 favors the spreading of tau via extracellular vesicles. Sci. Rep. 9, 9477 (2019).

58. Deming, Y. et al. The MS4A gene cluster is a key modulator of soluble TREM2 and alzheimer’s disease risk. Sci. Transl. Med. 11, (2019).

59. Werling, D. M., Parikshak, N. N. & Geschwind, D. H. Gene expression in human brain implicates sexually dimorphic pathways in autism spectrum disorders. Nat. Commun. 7, 10717 (2016).

60. Krabbe, G. et al. Microglial NFκB-TNFα hyperactivation induces obsessive-compulsive behavior in mouse models of progranulin-deficient frontotemporal dementia. Proc. Natl. Acad. Sci. U. S. A. 114, 5029–5034 (2017).

61. Li, B. & Dewey, C. N. RSEM: Accurate transcript quantification from RNA-Seq data with or without a reference genome. BMC Bioinformatics 12, 323 (2011).

62. Robinson, M. D. & Oshlack, A. A scaling normalization method for differential expression analysis of RNA-seq data. Genome Biol. 11, R25 (2010).

63. Robinson, M. D. & Smyth, G. K. Small-sample estimation of negative binomial dispersion, with applications to SAGE data. Biostatistics 9, 321–332 (2008).

64. Love, M. I., Huber, W. & Anders, S. Moderated estimation of fold change and dispersion for RNA-seqN data with DESeq2. Genome Biol. 15, 1–21 (2014).

65. Carpenter, B. et al. Stan: A probabilistic programming language. J. Stat. Softw. 76, (2017).

66. The 1000 Genomes Project Consortium et al. A global reference for human genetic variation. Nature 526, 68–74 (2015).

67. Delaneau, O., Marchini, J. & Zagury, J.-F. A linear complexity phasing method for thousands of genomes. Nat. Methods 9, 179–181 (2012).

68. Howie, B., Fuchsberger, C., Stephens, M., Marchini, J. & Abecasis, G. R. Fast and accurate genotype imputation in genome-wide association studies through pre-phasing. Nat. Genet. 44, 955–959 (2012).

69. Das, S. et al. Next-generation genotype imputation service and methods. Nat. Genet. 48, 1284–1287 (2016).

70. McCarthy, S. et al. A reference panel of 64,976 haplotypes for genotype imputation. Nat. Genet. 48, 1279–1283 (2016).

71. Chatterjee, N., Shi, J. & García-Closas, M. Developing and evaluating polygenic risk prediction models for stratified disease prevention. Nat. Rev. Genet. 17, 392–406 (2016).

72. Mitchell, T. J. & Beauchamp, J. J. Bayesian variable selection in linear regression. J. Am. Stat. Assoc. 83, 1023–1032 (1988).

73. Westra, H.-J. et al. Cell Specific eQTL Analysis without Sorting Cells. PLoS Genet. 11, e1005223.EP (2015).

74. VanderWeele, T. J., Mumford, S. L. & Schisterman, E. F. Conditioning on intermediates in perinatal epidemiology. Epidemiology 23, 1–9 (2012).

